# Timing variability and midfrontal ~4 Hz rhythms correlate with cognition in Parkinson’s disease

**DOI:** 10.1101/2020.10.26.356154

**Authors:** Arun Singh, Rachel C. Cole, Arturo I. Espinoza, Aron Evans, Scarlett Cao, James F. Cavanagh, Nandakumar S. Narayanan

## Abstract

Patients with Parkinson’s disease (PD) can have significant cognitive dysfunction; however, the mechanisms for these cognitive symptoms are unknown. Here, we used scalp electroencephalography (EEG) to investigate the cortical basis for PD-related cognitive impairments during interval timing, which requires participants to estimate temporal intervals of several seconds. Time estimation is an ideal task demand for investigating cognition in PD because it is simple, requires medial frontal cortical areas, and recruits basic executive processes such as working memory and attention.

However, interval timing has never been systematically studied in PD patients with cognitive impairments. We report three main findings. First, 71 PD patients had increased temporal variability compared to 37 demographically-matched controls, and this variability correlated with cognitive dysfunction as measured by the Montreal Cognitive Assessment (MOCA). Second, PD patients had attenuated ~4 Hz EEG oscillatory activity at midfrontal electrodes in response to the interval-onset cue, which was also predictive of MOCA. Finally, trial-by-trial linear mixed-effects modeling demonstrated that cue-triggered ~4 Hz power predicted subsequent temporal estimates as a function of PD and MOCA. Our data suggest that impaired cue-evoked midfrontal ~4 Hz activity predicts increased timing variability that is indicative of cognitive dysfunction in PD. These findings link PD-related cognitive dysfunction with cortical mechanisms of cognitive control, which could advance novel biomarkers and neuromodulation for PD.

## INTRODUCTION

Parkinson’s disease (PD) is a devastating neurodegenerative disease that involves motor as well as cognitive symptoms^1–4^. For motor symptoms of PD, detailed neurophysiological studies led to highly targeted and effective therapies such as deep-brain stimulation (DBS)^5–8^. Compared to motor symptoms, neurobehavioral symptoms are less treatable and have a stronger impact on quality of life^9, 10^. There are no therapies that improve cognitive symptoms of PD, in part because the neural mechanisms are unknown.

Cognitive deficits in PD are diverse, affecting a range of executive processes such as working memory, attention, reasoning, visuospatial dysfunction, inhibitory control, and flexibility^11, 12^. One process that is ideal for studying PD-related cognitive dysfunction is interval timing, which requires participants to estimate temporal intervals of several seconds^13^. Interval timing involves cortical areas that are dysfunctional in PD, such as the medial frontal cortex^14, 15^. Moreover, interval timing depends on executive functions such as working memory for temporal rules and attention to the passage of time^16^. PD patients have impairments in interval timing^17, 18^ and temporal processing in general^19, 20^. Since this prior work has focused on basal ganglia mechanisms in cognitively high-functioning PD patients it remains unknown how interval timing is affected by a broader range of cognitive impairments.

Findings from scalp electroencephalography (EEG) has provided convergent evidence of impaired midfrontal ~4 Hz rhythms in PD patients, particularly during interval timing^21, 22^. These cortical rhythms originate from medial frontal brain structures and are highly similar to midfrontal ~4-8 Hz rhythms that have been advanced as a mechanism of cognitive control^23–25^. Animal studies have shown that ~4 Hz oscillations are coherent with single neurons involved in the intricacies of cognitive operations^21, 22, 25, 26^. Midfrontal ~4 Hz rhythms are consistently impaired in PD patients, particularly around imperative cues signifying surprise, conflict, or the start of time estimation during interval timing tasks^27, 28^. Together, this line of evidence suggests that PD patients have dysfunctional cognitive control processes that can be indexed by ~4 Hz midfrontal EEG signals. This hypothesis specifically predicts that interval timing performance should be related to midfrontal ~4 Hz rhythms, and to cognitive dysfunction in PD. We directly tested this hypothesis by collecting EEG data from a large number of PD patients who demonstrated a range of cognitive abilities during the performance of an interval timing task. We used a time-production version of interval timing that we could relate to previous studies in PD and prior animal work^13, 17, 18, 21, 22, 29^. We found that the average temporal estimates of PD patients and demographically-matched controls were similar, yet variance was increased in PD. Critically, we discovered that this variability correlated with cognitive function in PD patients. We also found that PD patients had attenuated midfrontal cue-triggered ~4 Hz power relative to controls, and this neural measure also correlated with cognitive dysfunction in PD. Trial-by-trial analyses demonstrated that cue-related ~4 Hz power predicted timing variability. These findings support a model that cognitive failures in PD are related to attenuated cortical cognitive-control mechanisms, and that the integrity of this process can be monitored by EEG assessment of ~4 Hz rhythms.

## RESULTS

We tested the hypothesis that interval timing performance is related to cognitive dysfunction in PD. We recruited PD patients and demographically-matched controls to perform a two-interval-timing task with 3- and 7-second intervals. Participants started estimating when an imperative “Go” cue appeared in the center of their computer screen (Fig. 1a). Participants were instructed to press the keyboard spacebar just before they estimated the interval to have elapsed; we focused on this keypress as the start of their temporal estimate (Fig. 1b-c). Consistent with past work, average keypress time was similar for PD patients and demographically-matched controls for 3-second and 7-second intervals (mean+/− SEM keypress: Control 3-second: 3.2+/−0.1s, Control 7-second: 6.1+/−0.2s; PD 3-second: 3.2+/−0.1s, PD 7-second: 6.4+/−0.2s; Fig. 1d-e)^17, 18,30^. PD patients had more variance in keypress time as measured by the coefficient of variation (CV) for both the 3-second (Control: 0.19+/−0.01; PD: 0.23+/−0.01s; t(81)=-2.30, p=0.02; *d*=-0.52; Fig. 1f) and 7-second (Control: 0.15+/−0.01s; PD: 0.22+/−0.01s; t(106)=-3.24, p=0.002; *d*=-0.66; Fig. 1g) intervals. Notably, 3-second and 7-second CVs were similar (rho=0.79; p<0.0000001). Linear mixed-effects modeling revealed that there were main effects of interval-type (F(1,110)=5.4, p=0.02) and group (F(1,157)=28.2, p=0.0000004), but no significant interactions. Because keypress CV is a metric of temporal variability, these data provide evidence that PD affects the variability of temporal estimates^31, 32^.

**Fig. 1.**
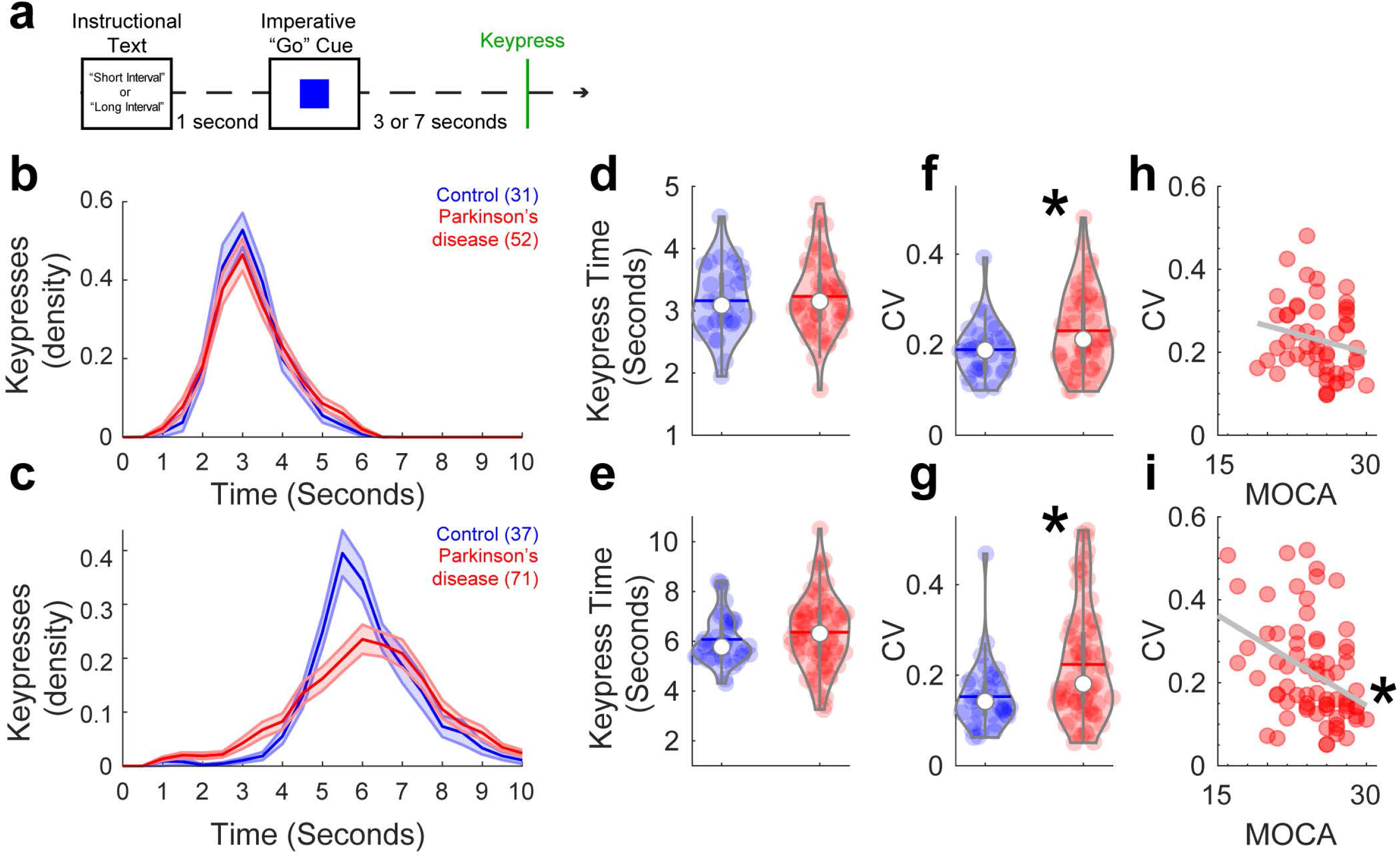
Timing variability correlated with cognitive dysfunction in PD. a) PD patients with a range of cognitive function and demographically-matched controls performed an interval-timing task at one of two intervals: 3 seconds or 7 seconds. This task required participants to press the spacebar (termed the “keypress”) when they estimated the target interval to have elapsed. b) Density estimates of keypress over time during 3-second intervals. Of recruited participants, only 31 controls and 52 PD patients had enough trials to analyze for 3-second intervals. c) Density estimates of keypress over time for 7-second intervals from controls (n=37) and PD (n=71) patients. Mean response times for d) 3-second and e) 7-second intervals for control (blue) and PD patients (red). Timing variability as measured by the coefficient of variation (CV) of keypress for f) 3-second and g) 7-second trials; *=p<0.05 via a t-test. Keypress CV correlated with cognitive function as measured by MOCA for h) 3-second and i) 7-second intervals. *=p<0.05 via Spearman’s rho.

Critically, we wanted to understand whether temporal variability was related to cognitive function as measured by MOCA, a widely-used clinical assessment of cognitive function that does not involve temporal judgement. In line with our hypothesis, we found that for 7-second intervals, cognitive dysfunction predicted keypress CV for PD patients (Spearman’s rho=-0.39, p=0.001; Fig. 1i). For 3-second intervals, there was no reliable correlation between MOCA and CV for PD patients (rho=-0.22; p=0.12; Fig.1h) although Fisher’s z comparisons with 7-second intervals were not statistically significant. However, 7-second CV correlations with MOCA were stronger in PD patients than controls, in whom there was no relationship of CV with MOCA (CV and MOCA; 3-second: rho=0.12, p=0.51; 7-second: rho=0.18, p=0.27; Fig. S1. Fisher’s z comparing CV and MOCA correlation between PD and controls, p<0.004). For 3-second intervals in PD patients, there was no significant relationship of keypress CV with mUPDRS (rho=-0.08, p=0.59; Fig. S2a). In PD patients, keypress CV for 7-second intervals was also correlated with mUPDRS (rho=0.28; p=0.02; Fig. S2b), though MOCA and mUPDRS were not correlated (rho=-0.09, p=0.47; Fig. S2c). Hierarchical multiple linear regression analysis supported the relationship between CV and mUPDRS (F(1,69)=4.3; p=0.04). However, the model became much stronger when MOCA was added after mUPDRS in the regression analysis (F(2,68)=7.6; p=0.001), and regression coefficients differed significantly between the two models (F(1,68)=10.3; p=0.002). These results suggest that there was a significant effect of MOCA above and beyond the mUPDRS in PD patients during the 7-second interval timing task. Taken together, these data support our hypothesis that temporal variability as measured by keypress CV was linked to cognitive dysfunction in PD.

To explore the neural basis of these deficits, we analyzed EEG data. We focused on two key events: the imperative “Go” cue and the keypress. In addition, we focused on the midfrontal electrode Cz in line with our prior work^21, 22, 27–29, 33^. Power-spectral density (PSD) analyses of resting-state data is shown in Figure S3. A comparison of ERPs or CNV did not reveal consistent differences between control and PD patients (Figs. S4–6). In our time-frequency analysis, we found that PD patients had attenuated midfrontal delta rhythms triggered by the imperative cue (300-400 ms after cue, 1-4 Hz Control: 0.7+/−0.2 dB, PD: 0.0+/−0.1 dB; t(106)=2.31, p=0.02; *d*=0.47 for 7-second trials; 3-second trials in Fig.S7; midfrontal cluster of electrodes FCz/Cz/CPz in Fig.S8). We also found that PD patients had attenuated theta rhythms (300-400 ms after cue; 4-7 Hz Control: 0.3+/−0.2 dB, PD: −0.3+/−0.1 dB; t(106)=2.24, p=0.03; d=0.45; Fig. 2a-c, d and f).

**Fig. 2.**
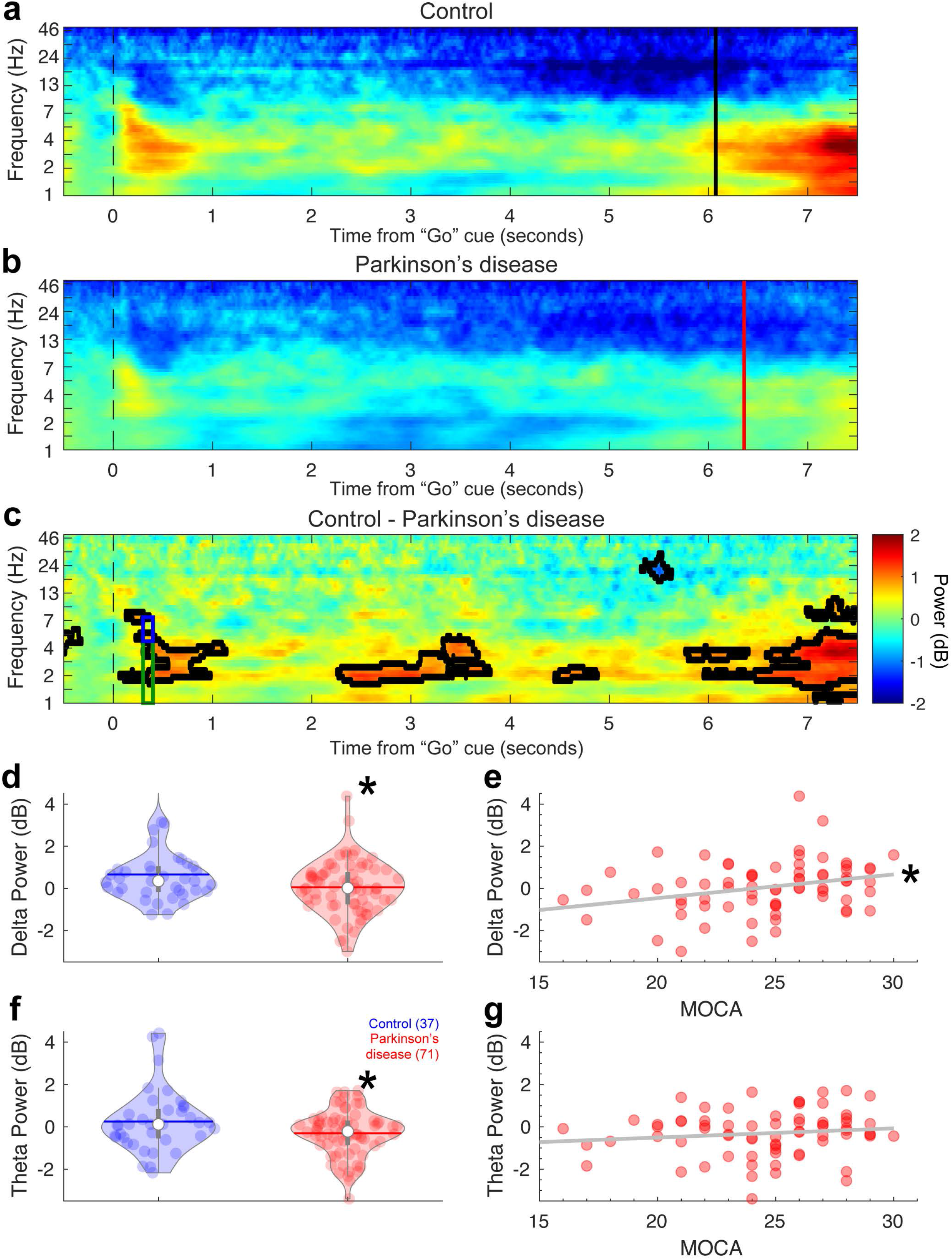
Cue-triggered midfrontal delta power predicts cognitive dysfunction in PD. Frequency power of midfrontal activity over time from the imperative “Go” cue from a) control and b) PD participants on 7-second trials from EEG electrode Cz. c) Comparison of control and PD patients. Areas outlined by solid black lines indicate p<0.05 via a t-test of activity in control compared to PD participants. There was significantly less d) cue-triggered midfrontal delta power (1-4 Hz, time-frequency-Region-of-interest (tf-ROI): green box) and f) cue-triggered midfrontal theta power (4-7 Hz, tf-ROI: blue box) in controls vs. PD patients. e) Delta power predicted cognitive dysfunction as measured by MOCA in PD patients, but g) theta power did not. *=p<0.05. Data from control (n=37) and PD (n=71) patients.

Phase-locking analyses revealed that PD patients had marked differences in phase-locking around ~4 Hz on 7-second trials (Fig.S9). Attenuated cue-triggered delta activity replicates prior work from our group, showing that PD patients had disrupted ~4 Hz activity around events that engage cognitive control such as novelty, conflict, and imperative timing cues^21, 22, 27–29^. As in our prior work, distinctions between ~4 Hz activity were more reliable for the longer 7-second intervals than for 3-second intervals (Fig. S7) or for keypress (Fig. S10)^21, 22^.

Strikingly, we found that midfrontal cue-triggered delta rhythms in the 7-second task correlated with cognitive dysfunction in PD (rho=0.31, p=0.01; Fig. 2e; Fig. S11). This was not observed for theta rhythms (rho=0.15, p=0.21; Fig. 2g), although Fisher’s z test did not reveal reliable differences between delta and theta correlations with MOCA. Delta power was correlated with CV (rho=-0.4, p=0.006), and mediation analysis confirmed that CV significantly mediated the relationship between delta power and MOCA (Average Causal Mediation Effects: p<0.05), but mUPDRS did not (p>0.05). This finding was directly supportive of our hypothesis that deficits in midfrontal cue-triggered ~4 Hz rhythms underlie cognitive deficits in PD.

These data suggest that PD patients have attenuated cue-triggered delta rhythms, and that individuals with smaller delta activity have larger CV and lower MOCA scores. To test the hypothesis that cue-triggered delta-dysfunction mechanistically affects time estimation ~7 seconds later, we turned to linear mixed-effects models at a trial-by-trial level. To address this trial-by-trial hypothesis, we turned to linear mixed-effects models. We used a model where the outcome variable was keypress as an index of time estimation, and predictor variables were EEG delta power (1-4 Hz, 300-400 ms after cue), interval (short vs. long), disease status (control vs. PD), and cognitive status (as measured by MOCA). As previously shown, this analysis revealed the main effect of short vs. long intervals (Table 2) and a significant interaction between delta power and disease status (F(1,6421)=4.48, p=0.03; Table 2). In support of our trial-to-trial hypothesis, there was an interaction between delta power, disease status, and MOCA (F(1,6421)=4.71, p=0.03; Table 2). There were no higher interactions with interval.

Visualization of these relationships revealed that for controls, delta power was inversely related to keypress variance at a trial-specific level, such that lower delta power was associated with more variable keypress around the 7-second interval (Fig. 3a). However, for PD patients who had lower overall delta power there was no clear relationship between delta power and keypress (Fig. 3b; see Fig. S12 for 3-second intervals). These data linking cue-triggered midfrontal delta power and keypress support a model whereby in control participants, normative cue-triggered delta activity relates to better precision of time-estimates, yet this relationship is absent in the PD group.

**Fig. 3.**
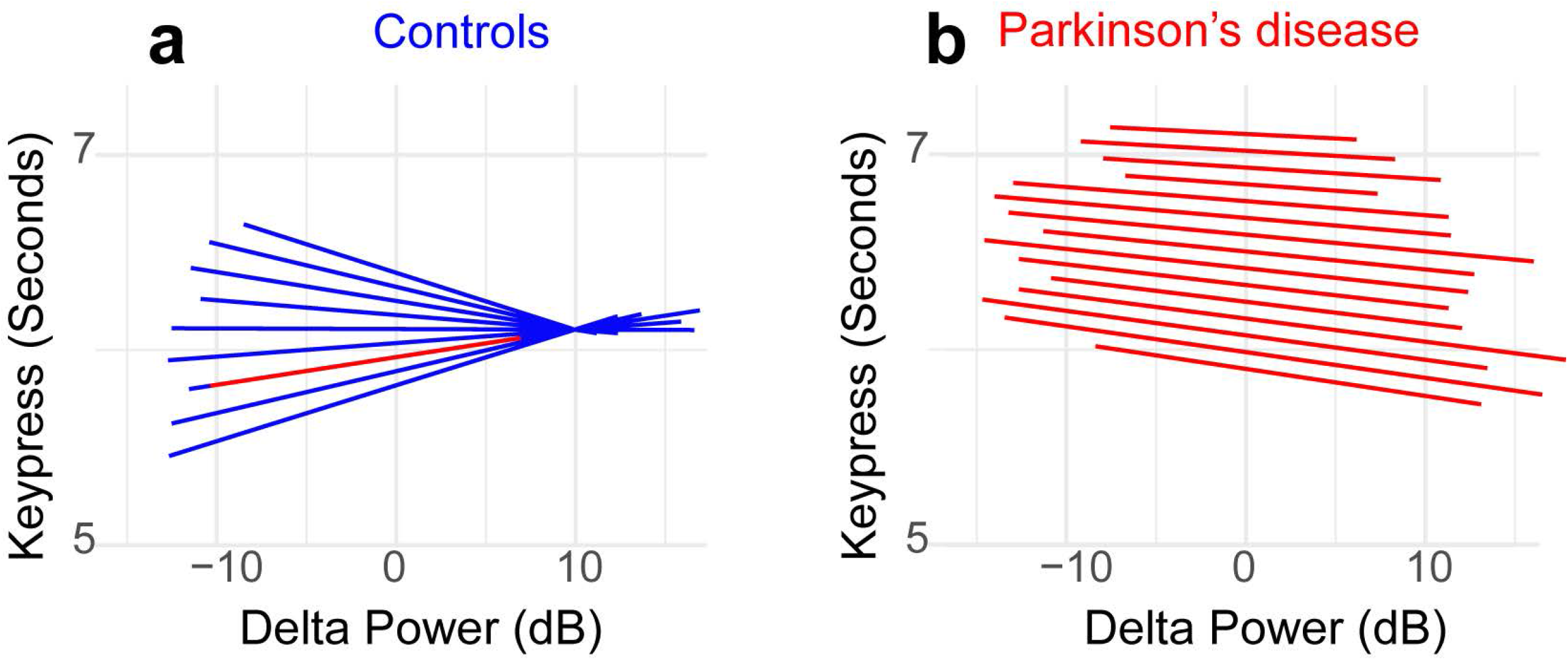
Increased cue-triggered delta activity predicts decreased temporal variability in controls but not in PD. We plotted the slopes for each individual from our trial-by-trial linear mixed-effects model trial-by-trial delta power on keypress. This analysis revealed that for a) Controls, increased midfrontal delta-power was associated with keypress that was more clustered around ~7 seconds; individual slopes plotted as blue lines. However, for b) PD patients, there was no clear relationship between delta power and keypress; individual slopes plotted as red lines. These data indicate that for controls, increased delta power was linked with less timing variability, while PD patients (who had lower delta power overall – see Fig 2b) had higher timing variability and less of an influence between delta power and performance. Data from all trials in control (n=37) and PD (n=71) patients.

Finally, we examined the effects of levodopa on ~4 Hz rhythms during interval timing. We found that levodopa did not reliably alter keypress time (Fig. S13a). However, it did tighten CVs for 3-second intervals (ON: 0.20+/−0.03, OFF: 0.16+/−0.02; paired t(8)=2.75, p=0.03; Fig. S13b) though not for 7-second intervals (ON: 0.18+/−0.03, OFF: 0.15+/−0.02; paired t(8)=1.83, p=0.11; Fig. S13b). Critically, there were no reliable effects of levodopa on cue-triggered midfrontal delta/theta rhythms (Fig. S13c-e). This finding is in line with our past work and consistent with the lack of levodopa’s effects on cortical circuits^28, 34^.

## DISCUSSION

This study revealed that deficits in midfrontal cognitive control processes contribute to increased variability in interval timing and to cognitive dysfunction in PD. First, we found that PD patients had increased timing variability that correlated with cognitive function. Second, we replicated our earlier work demonstrating that PD patients had deficits in cue-triggered midfrontal delta activity^21, 22, 27, 28^ and discovered that these deficits also correlated with cognitive dysfunction in PD. Finally, we found that delta activity was linked with cognitive function during interval-timing performance in PD at a trial-by-trial level. Our data illuminate the nature of timing deficits in PD and PD-related cognitive dysfunction. Indeed, these results suggest that increased timing variability is a manifest expression of dysfunctional midfrontal ~4 Hz rhythms which contribute to cognitive dysfunction in PD. Together, these interindividual and intraindividual findings suggest a candidate mechanism and biomarker of cognitive dysfunction PD.

Complex cellular factors such as alpha-synuclein and deficits in ascending neurotransmitter systems contribute to PD^11, 35^. Despite this complexity, these cellular processes lead to neurophysiological phenomena observable by scalp EEG and intracortical recordings, which have been enormously helpful in both designing medical therapies and targeting brain stimulation sites to address the motor symptoms of PD^33, 36–38^. Our results are significant in advancing our neurophysiological understanding of cognitive function in PD because we link PD-related cognitive dysfunction (as measured by MOCA) to midfrontal ~4 Hz rhythms, which are a putative cognitive control mechanism^23^. These findings imply that cognitive dysfunction in PD patients can result in part from their inability to recruit cognitive control processes. Our data support a model in which deficits in ~4 Hz cognitive control signals contribute to cognitive dysfunction in PD patients.

Our work also provides insight into the nature of timing deficits in PD patients. Seminal work in cognitively normal PD patients described deficits in working memory processes during interval timing as a function of dopamine signaling^17, 18^. Past work has linked PD-related timing deficits to basal-ganglia dysfunction^13, 19^; by contrast, our work strongly implicates cortical cognitive control mechanisms. In our study, some cognitively impaired PD patients did not reliably release the spacebar. Thus, our primary behavioral measure was keypress, or the start of when participants had estimated intervals to have elapsed. Hence our results are not directly comparable to the peak-time metrics in past studies. Of note, prior work on interval-timing in PD has focused on cognitively high-functioning patients^19, 20^. We included the full-range of cognitive function in PD, which is likely more reflective of real-world PD but may limit behavioral assessments^1, 3, 4^. We found that PD patients have a high rate of variability in keypress that was proportional to their cognitive impairment. Correlations between timing variability and cognition, and differences in delta activity for PD patients were more reliable at the 7-second interval compared to the 3-second interval, although we note that fewer patients performed the 3-second interval^22, 29^. This may be a result of distinct motor mechanisms affecting keypress at 3-second or increased working-memory / cognitive load for 7-second intervals. Furthermore, our interval-timing task included a strong motor component in key press, and some studies of interval timing that use non-motor tasks have failed to find differences^39^. These data, combined with our differences at 3 vs. 7 seconds, may support a view where sustained attention to the passage of time is altered in PD.

Sustained attention is necessary for time estimation and may reflect facets of cognitive control processes that malfunction in PD patients with cognitive dysfunction. Low-frequency rhythms are common to the effortful exertion underlying cognitive-control processes related to error-processing, conflict resolution, novelty detection, as well as working memory^23, 40^. Impaired cognitive control in PD patients may be linked to decreased 4 Hz rhythms and greater interval-timing variability^23^. Additionally, our task included distractors which involve additional executive function, and it is possible that this preferentially affected PD patients with cognitive dysfunction. We have found that cortical neurons involved in temporal processing can be coherent at ~4 Hz^21, 25, 41, 42^.

Indeed, our data show that a cue-triggered burst of ~4 Hz activity engages neurons that are directly involved in temporal processing^22, 25, 41, 42^. Such low-frequency oscillations can engage not only cortical neurons but also neurons in the subthalamic nucleus or other brain areas^29, 43, 44^. Our data suggest that in cognitively limited PD patients, delta power is decreased and results in inefficient engagement of temporal processing by frontostriatal networks^45–47^. If neuronal ensembles encoding time are not precisely engaged, they might lead to decreased precision in time estimates several seconds later^47^. Our findings that delta power predicts keypress variability supports this model.

These findings are in line with recent evidence that ~4 Hz stimulation improves cognitive function in PD^29, 48^. Animal studies support this idea; indeed, highly specific stimulation at low frequencies can improve interval timing in animal PD models^21, 41, 49^ and other cognitive disorders such as schizophrenia^49^. Strikingly, human studies show that ~4 Hz subthalamic nucleus DBS can improve interval timing and conflict tasks, yet it remains unknown what effect cortical stimulation might have. Low-frequency subthalamic nucleus DBS has the potential to mitigate cognitive control deficits in PD, but precise and extensive circuit-mapping will be required to determine whether low-frequency brain stimulation can improve real-world function in PD patients. This effort will also require systematic clinical studies using EEG-based biomarkers to assess the usefulness of ~4 Hz rhythms in diagnosing cognitive dysfunction, which is often under-recognized in the clinical setting^50^. Of note, we focus on ~4 Hz based on extensive human work in PD and cognitive control as well as our animal work^21–23, 25, 28, 33^. While ~4 Hz deficits may not be highly predictive of PD, our work suggests that interval-timing variability and ~4 Hz deficits are predictive of cognition in PD.

Midfrontal electrodes are reflective of medial frontal generators^29^, yet many studies of timing thus far have focused on basal-ganglia processes^19^. Levodopa strongly affects nigrostriatal dopamine, resulting in shifts in time estimation^17, 18^. However, levodopa does not reliably predict cortical dysfunction or cognitive deficits in PD^35, 51, 52^. PD can affect dopamine signaling in the cortex, and our work clearly links increased timing variability to cortical ~4 Hz rhythms. Although levodopa can affect frontal circuits in complex ways^52, 53^, our data here are consistent with prior findings that delta/theta rhythms are not sensitive to levodopa^28, 54, 55^. Of note, our sample size was small, and practice effects may be relevant as patients performed ‘OFF’ sessions after ‘ON’, and a prior study showed that levodopa alone can affect interval timing^28^. However, we are not aware of any work demonstrating that levodopa can affect cortical 4 Hz rhythms^20, 56^. Identifying the defects in cognitive control mechanisms that occur in the midfrontal cortex and are defective in PD may be helpful in designing therapies for PD which target these circuits.

Our work has several limitations. First, scalp EEG has poor spatial resolution, and more advanced MEG, EEG/fMRI, or intraoperative recording studies might make it possible to draw more detailed inferences about the sources of altered temporal processing in PD. Second, it is possible that neuropsychological assays more advanced than MOCA will further clarify advanced cognitive or interval timing deficits in PD. One promising feature of interval timing is that it is highly translatable to animal models, enabling detailed insights into cortical mechanisms of temporal control of action. Third, our task involves a strong motor component. However, mediation analyses suggested that motor function in PD as measured by the mUPDRS was not related to timing-variability or cognition (Fig. S2), and neither was response-related activity (Fig. S10).

Fourth, we presented distractors, but we lack vocal recordings to analyze these events. Fifth, we did not observe strong beta-rhythm distinctions in PD patients (Figs. S3 and S10); however, our PD population is considerably more diverse than prior studies, and beta rhythms can be sensitive to movement, which may have undermined our ability to detect these differences. Finally, although our work here is correlative, it will guide future studies aimed at establishing the cellular and network mechanisms contributing to cognitive dysfunction in PD.

In summary, this study provides novel evidence that PD patients have increased variability in interval-timing performance. Both increased timing variability and midfrontal delta activity correlated with cognitive dysfunction in PD, and greater delta activity corresponded to more precise time estimates. This work suggests that impaired cognitive control mechanisms contribute to cognitive dysfunction in PD.

## METHODS

### Participants

Eighty-nine PD patients (56 men and 33 women; Table 1) were recruited from the movement disorders clinic at the University of Iowa. All patients were examined by a movement-disorders physician to verify that they met the diagnostic criteria recommended by the United Kingdom PD Society Brain Bank criteria. Forty-one control participants were recruited from the Iowa City community and matched for age, sex, and education. All PD and control participants were determined to have the decisional capacity to provide informed consent in accordance with the Declaration of Helsinki and the Ethics Committee on Human Research. We obtained written informed consent from each participant. Eighty PD patients participated in the study while taking their prescribed medications, including levodopa, as usual. Because levodopa can affect interval timing^17, 18^, we also recruited nine additional PD patients for comparisons of interval timing and EEG signals on and off dopaminergic medications. We first tested these PD patients when they were taking medications as usual (ON sessions), and then they were asked to withhold dopaminergic medications for 12 hours (OFF sessions) prior to repeating behavioral and EEG testing. All research protocols were approved by the University of Iowa Human Subjects Review Board (IRB# 201707828).

**Table 1.** Demographic, disease, non-motor, motor, and cognitive characteristics.

|  | Control (N = 41) | PD (n = 80) | ON (n = 9) | OFF (n = 9) |
| --- | --- | --- | --- | --- |
| **Demographics and Disease** |  |  |  |  |
| Gender, M/F | 56/24 | 23/18 | 6/3 | 6/3 |
| Age, years | 71.3 (1.2) | 68.7 (0.9) | 67.2 (1.8) | 67.2(1.8) |
| Disease Duration, years | - | 5.2 (0.4) | 3.7 (1.2) | 3.7 (1.2) |
| LED, mg/day | - | 860.5 (50.3) | 727.8 (143.9) | - |
| **Cognition Characteristics** |  |  |  |  |
| MOCA (0-30) | 26.6 (0.3) | **24.3 (0.4)\*\*** | 27.1 (0.6) | 27.7 (0.7) |
| **Non-Motor Characteristics** |  |  |  |  |
| PDSS (0-60) | 7.2 (0.7) | **17.5 (1.1)\*\*** | 12.8 (2.3) | 12.8 (2.3) |
| BAI (0-63) | 1.8 (0.3) | **14.7 (1.2)\*\*** | 9.4 (2.8) | 9.4 (2.8) |
| GDS (0-15) | 0.7 (0.2) | **3.7 (0.4)\*\*** | 2.8 (0.7) | 2.8 (0.7) |
| **Motor Characteristics** |  |  |  |  |
| mUPDRS (0-56) | - | 13.4 (0.8) | 9.1 (1.6) | **11.6 (1.7)\*** |

Values are expressed as mean (standard error of mean). A Two-sample *t* test was used for comparison between PD vs control subjects. Paired *t* test was used for comparison between ON vs OFF dataset. Significant effects in bold. \*\*= $p < 0.01$ , \*= $p < 0.05$ .
Abbreviations: Male (M); Female (F); Montreal Cognitive Assessment (MOCA); Parkinson's Disease Sleep Scale (PDSS); Beck Anxiety Index (BAI); Geriatric Depression Scale (GDS); motor Unified Parkinson's Disease Rating Scale (mUPDRS).

**Table 2.** Linear mixed-effect model of interval, disease, delta activity and cognitive function on keypress. (R^2^=0.46)

| Contrast | DF | F | p |
| --- | --- | --- | --- |
| Delta | 6421 | 1.06 | 0.30 |
| CTL/PD | 102 | 0.12 | 0.73 |
| MOCA | 102 | 0.79 | 0.38 |
| Interval | 6446 | 134.29 | **<0.0000001\*\*** |
| Delta:CTL/PD | 6421 | 4.48 | **0.03\*** |
| Delta:MOCA | 6421 | 0.88 | 0.35 |
| CTL/PD:MOCA | 102 | 0.08 | 0.77 |
| Delta:Interval | 6416 | 1.42 | 0.23 |
| CTL/PD:Interval | 6446 | 1.20 | 0.27 |
| MOCA:Interval | 6447 | 19.11 | **0.00001\*\*** |
| Delta:CTL/PD:MOCA | 6421 | 4.71 | **0.03\*** |
| Delta:CTL/PD:Interval | 6416 | 0.06 | 0.81 |
| Delta:MOCA:Interval | 6416 | 1.40 | 0.24 |
| CTL/PD:MOCA:Interval | 6447 | 1.08 | 0.30 |
| Delta:CTL/PD:MOCA:Interval | 6416 | 0.14 | 0.71 |

Significant effects in bold via an F test in R. \*\*= $p<0.01$ , \*= $p<0.05$
Abbreviations: Degrees of Freedom (DF); Control (CTL), Parkinson's disease (PD), Montreal Cognitive Assessment (MOCA).

We analyzed data from those patients who completed our experimental protocol and whose midfrontal EEG channels did not produce immediately observable artifact or noise. Of 130 recruited participants, 6 PD patients and 4 controls were excluded from further analysis because they did not meet these criteria. Thus, EEG data from a total of 83 PD (74 ON + 9 ON/OFF) patients and 37 controls were available. Demographics of patients and control subjects are summarized in Table 1.

### Interval-timing task

All testing was done in a laboratory setting in the University of Iowa’s Department of Neurology. All stimuli were presented via Psychtoolbox v3.0, a MATLAB toolbox running on a Dell XPS workstation and a 19-inch monitor. We performed a peak-interval timing task with 3-second and 7-second randomly-intermixed intervals^17, 18, 57^. In this task, participants estimated the duration of either a 3-or 7-second interval after reading instructions displayed as text at the center of a video screen. White instructional text on the center of a black screen was presented that read “Short interval” on 3-second interval trials and “Long interval” on 7-second interval trials. The actual interval durations were never communicated to the patient. After the instructions were displayed for 1 second, the start of the interval was indicated by an imperative “Go” cue: the appearance of an image of a solid box in the center of the computer screen (Fig. 1a).

The cue was displayed on the screen for the entire trial, which lasted 8-10 seconds for 3-second intervals and 18-20 seconds for 7-second intervals. Participants were instructed to estimate the duration of the intervals without counting. As a distraction to discourage counting, a vowel appeared sporadically at random intervals in the center of the screen. Participants were instructed to press the keyboard spacebar at the start of when they judged the target interval to have elapsed, and to release the spacebar after they judged the target interval to have elapsed (Fig. 1a). Of note, our population included many patients with advanced motor dysfunction as well as cognitive impairment and dementia, and some patients did not reliably release the space bar once they had pressed it. Consequently, for all participants we focused our analyses only on the time participants pressed the space bar; this time is referred to as the “keypress.” On 15% of the trials, participants were then given nonnumerical/nonverbal performance feedback on the screen indicating their response relative to the actual target interval; feedback was a horizontal green line indicating how close their keypress was to the target time. Because feedback was relevantly infrequent, we did not analyze this event. The time between trials varied from 3-8 seconds. All participants performed 6 training trials prior to testing trials and verbalized understanding of the task. Testing sets consisted of 40 trials of each interval length, for a total of 80 trials. Trials were presented in a random order in blocks of 20 trials. 4 blocks were presented. Participants took a self-paced break between each block and moved to the next block by pressing any key. Only data from patients with >20 trials of each type of trial were analyzed; 71 PD patients and 37 controls had sufficient data on 7-second trials; of these, 52 PD patients and 31 controls had sufficient data on 3-second trials. No trials were removed based on EEG characteristics. We analyzed the mean keypress time to capture timing accuracy for PD patients. To analyze timing precision, we computed the coefficient of variation (CV) by dividing the standard deviation by the mean keypress time.

### EEG recording and analysis

Scalp EEG signals were collected from 64 channels of EEG actiCAP (Brain Products GmbH) with a 0.1 Hz high-pass filter and a sampling frequency of 500 Hz. Electrode Pz was used as a reference, and electrode Fpz was used as the ground. We used recording methods as described previously^28, 33^. We used a customized EEG cap with I1 and I2 leads and without PO3, and PO4, allowing for placement of cerebellar leads. Briefly, an additional channel was recorded at the mid-inion region (Iz) and we removed unreliable Fp1, Fp2, FT9, FT10, TP9, and TP10 channels, resulting in 59 channels for pre- and post-processing. Data were epoched around the onset of instructional text (−2 s to 10 s for 3-second intervals and −2 s to 20 s for 7-second interval), from which the associated imperative Go-cue-locked and corresponding responses-locked epochs were isolated.

EEG activity at the reference electrode Pz was recovered by computing the average reference. Bad channels and bad epochs were identified using the FASTER algorithm and pop_rejchan from EEGLAB, and were then interpolated and rejected respectively^58^. On average 1.6 +/−0.9 channels were removed, and Cz was never removed during preprocessing. Eye blinks were removed using independent component analysis (ICA). Since our *a priori* hypothesis was focused on midfrontal ~4 Hz activity, we based our analyses on the Cz vertex electrode in this report^21, 22, 27–29, 33^.

### Event-related potentials (ERPs)

ERPs were low-pass-filtered at 20 Hz. We quantified ERP differences by focusing on canonical events of negative or positive deflections in voltage. These ERP differences were then analyzed as the peak-to-trough difference in the dominant canonical cue-or response-locked morphological feature (e.g. P2 and N2 for cue, pre-response peak and error-related negativity (ERN) for response). Topographic maps were created to highlight the sensitivity of midfrontal areas to PD-related differences^59^. Contingent negative variation (CNV, which can be influenced by dopamine^60^), was calculated by computing mean amplitude from 500 – 3000 ms and 500 – 7000 ms around the “Go” cue during 3 s and 7 s interval timing tasks, respectively.

### Time-frequency analyses

We computed spectral measures by multiplying the fast Fourier transformed (FFT) power spectrum of single-trial EEG data with the FFT power spectrum of a set of complex Morlet wavelets (defined as a Gaussian-windowed complex sine wave: e^i2πtf^e^-t^2/(2xσ^2)^, where *t*=time and *f* =frequency). These wavelets increased from 1-50 Hz in 50 logarithmically-spaced steps. Steps defined the width or “cycles” of each frequency band, increasing from 3-10 cycles between 1-50 Hz and taking the inverse FFT^61^. The end result of this process was identical to time-domain signal convolution, and it resulted in estimates of instantaneous power (the magnitude of the analytic signal) and phase angle (the arctangent of the analytic signal). We then cut the length of each signal accordingly for each trial. As in our past work, cue-locked epochs lasted −500 – +7500 ms in the case of 7-second intervals, and −500 – +3500 in the case of 3-second intervals; note that these analyses were focused primarily on cue-locked events. For response-locked trials they were −500 – +1000 ms. These short temporal epochs reflect the wavelet-weighted influence of longer time and frequency periods. Power was normalized by converting to a decibel (dB) scale (10*log10(powert/powerbaseline)), allowing us to directly compare the effects across frequency bands. As in our past work, the baseline for each frequency was calculated by averaging power from ‑300 – −200 ms prior to the onset of the imperative “Go” cue^27, 28^. We note that specific frequencies can greatly vary between subjects, and frequency-band analyses can be diverse. Given that our primary hypothesis pertained to 1-4 Hz delta power following imperative cues, the time-frequency region of interest (tf-ROI) analysis was constrained to pre-defined frequency bands with the potential to be relevant (as determined by extensive past research). These bands included delta: 1-4 Hz, theta: 4-7 Hz, alpha: 8-12 Hz, and beta: 13-30 Hz^21–25, 27–29, 33^. We selected the cue-triggered 300-400 ms ROI a priori based on this prior work. In addition to this well-motivated tf-ROI, we used a cluster size of 500 pixels and a t-test as in our past work^27, 33^. We restricted all analyses to electrode Cz to be consistent with our prior work.

### Clinical Metrics

All study participants completed the Montreal Cognitive Assessment (MOCA), a cognitive screening tool for the evaluation of a variety of rudimentary cognitive assessments that do not involve temporal judgement (e.g., visuospatial ability, naming, short-term/working memory, etc.). Each item on the test is tabulated to give a total score ranging from 0-30, with lower scores indicating poorer cognitive function^62, 63^. Of our sample, 5 patients had dementia, 26 patients had mild-cognitive impairment, and 40 patients were normal^63^.

We assessed motor function using Part 3 of the Unified Parkinson’s Disease Rating Scale (mUPDRS)^64^. In addition, we performed other evaluations such as the PD sleep scale, the Geriatric Depression Scale, the Beck Anxiety Inventory, elements from the NIH Toolbox, the full UPDRS, and gait analyses in some patients, although these were not related to our hypothesis and were not analyzed.

### Statistical Analyses

Since our hypothesis pertained to interval timing performance and midfrontal ~4 Hz power between two groups (PD vs. control), we used t-tests to compare behavioral data between groups and for EEG analyses between groups, in line with our prior work^21, 22, 27–29^. Effect sizes were calculated via Cohen’s *d*, and relationships among behavioral measures, EEG features, and clinical metrics (such as MOCA and mUPDRS) were calculated using the non-parametric Spearman’s rho correlation. Correlations were compared by Fisher’s r to z transformation. EEG preprocessing was performed in the Matlab toolbox EEGLAB. Behavioral and EEG analyses were performed using custom-written scripts in MATLAB that were consistent with prior work**^28^**.

We performed hierarchical multiple linear regression in order to compare successive regression models in R statistical software, using the CV of response time. We tested both mUPDRS and MOCA score variables to determine whether or not adding MOCA after mUPDRS enhances predictive capability. These predictor models were compared using an ANOVA. We performed mediation analysis using the *mediation* package in R, which provides an estimate of the average causal mediation effects (ACME). P-values for these analyses were derived from 1000 iterations of bootstrapping, and they were expressed relative to alpha of p=0.05. Variables included in mediation analyses included “Go”-cue-locked power values, CV of response time, and MOCA score.

To construct a comprehensive model of time estimation on a trial-by-trial basis, we used linear mixed-effects modeling of every trial from all participants using the *lmer* package in R. We quantified trial-by-trial effects of cue-related EEG power on time estimation by constructing a linear mixed-effects model, where response time was the outcome variable. EEG power, disease status, MOCA score, and interval-type were predictor variables. The trial-by-trial relationship between response time and cue-related EEG power was computed and participant was included as a random effect to reduce bias in our modeling results by accounting for the inherent correlation that exists between repeated measurements on the same subject. Our model was built based on our hypothesis; model-comparison via Akaikie Information Criteria (functions *AIC* and *step* in R) revealed that our model had less prediction error than individual covariates (AIC for our model: 111639; AIC for delta only: 117168; PD only: 117157; Interval only: 111745; and MOCA only: 117155).

All data and code is available at narayanan.lab.uiowa.edu and PRED+CT (predict.cs.unm.edu)^65^.

## DATA AVAILABILITY

Data and code will be available on publication at PRED+CT and narayanan.lab.uiowa.edu.

## ACKNOWLEDGEMENTS

We would like to thank Matthew Mattell for feedback on drafts of this manuscript. This work was funded by R01NS100849-01A1 and the Roy J. Carver Charitable Trust.

## AUTHOR CONTRIBUTIONS

AS, AIE, RC, and NN participated in design of the study. AS, RC, SC, and AIE, and AE collected data for this manuscript. AS and NN analyzed the data, while RC independently checked analysis code. AS, RC, JC, and NN wrote the manuscript.

## COMPETING INTERESTS

The authors report no competing interests.

## ADDITIONAL INFORMATION

Additional Supporting Information is available in the online version of this article.

## Supplementary Figures

**Fig. S1.**
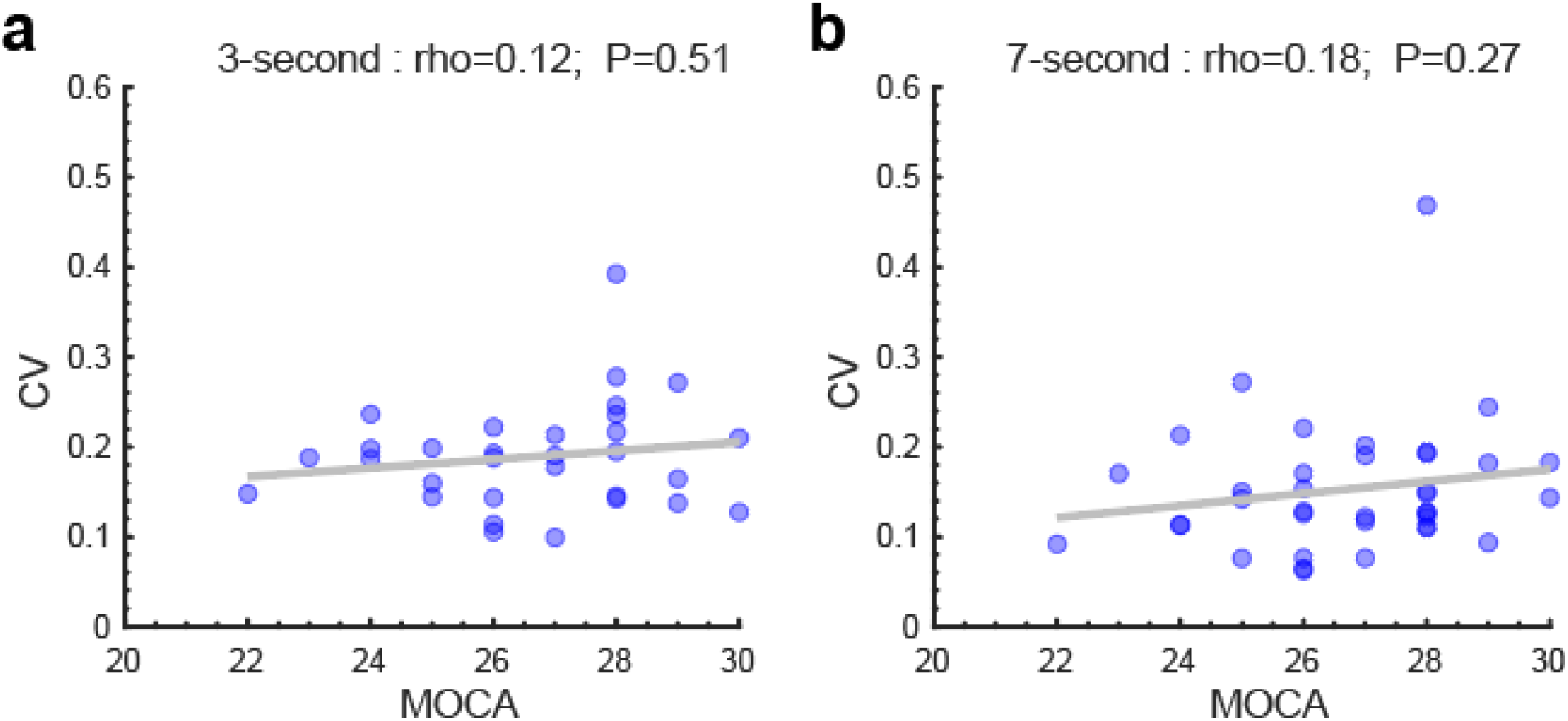
Timing variability does not correlate with cognitive dysfunction in controls. Timing variability as measured by keypress CV correlated with cognitive function as measured by MOCA for a) 3-second and b) 7-second intervals in control participants only.

**Fig. S2.**
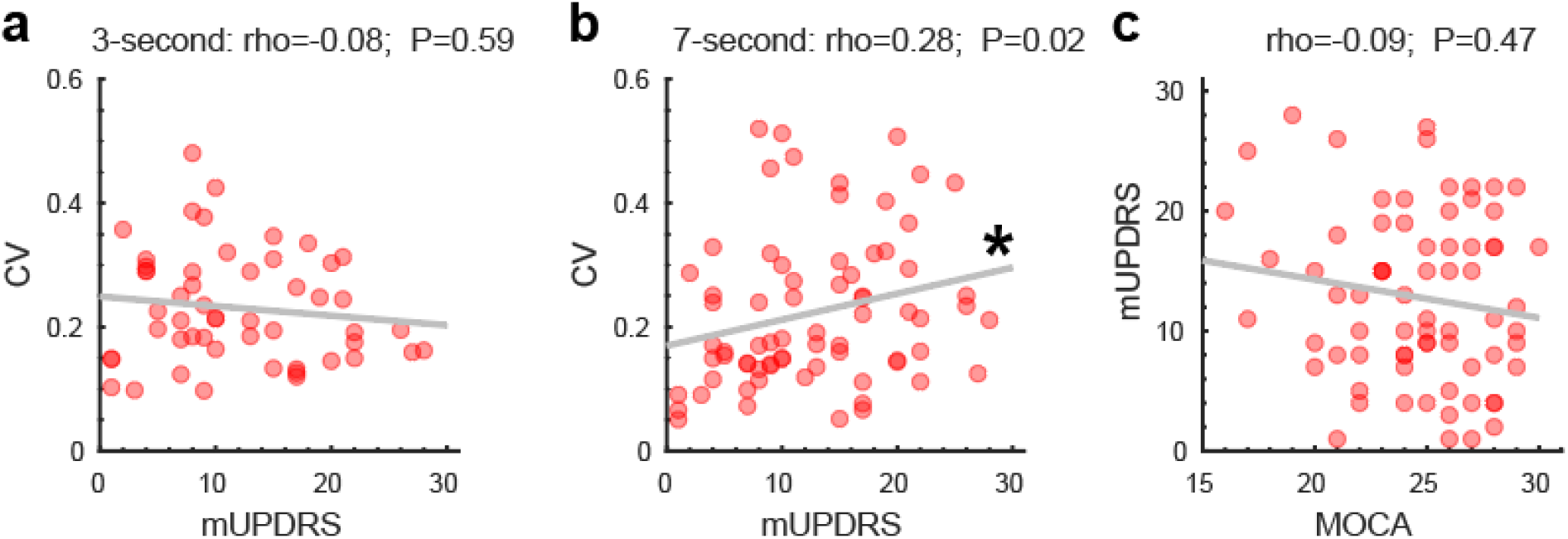
Motor function by mUPDRS correlated with CV and MOCA in PD. Keypress CV correlated with motor function as measured by the mUPDRS for a) 3-second and b) 7-second intervals (*=p<0.05). c) mUPDRS was not correlated with cognitive function as measured by MOCA.

**Fig. S3.**
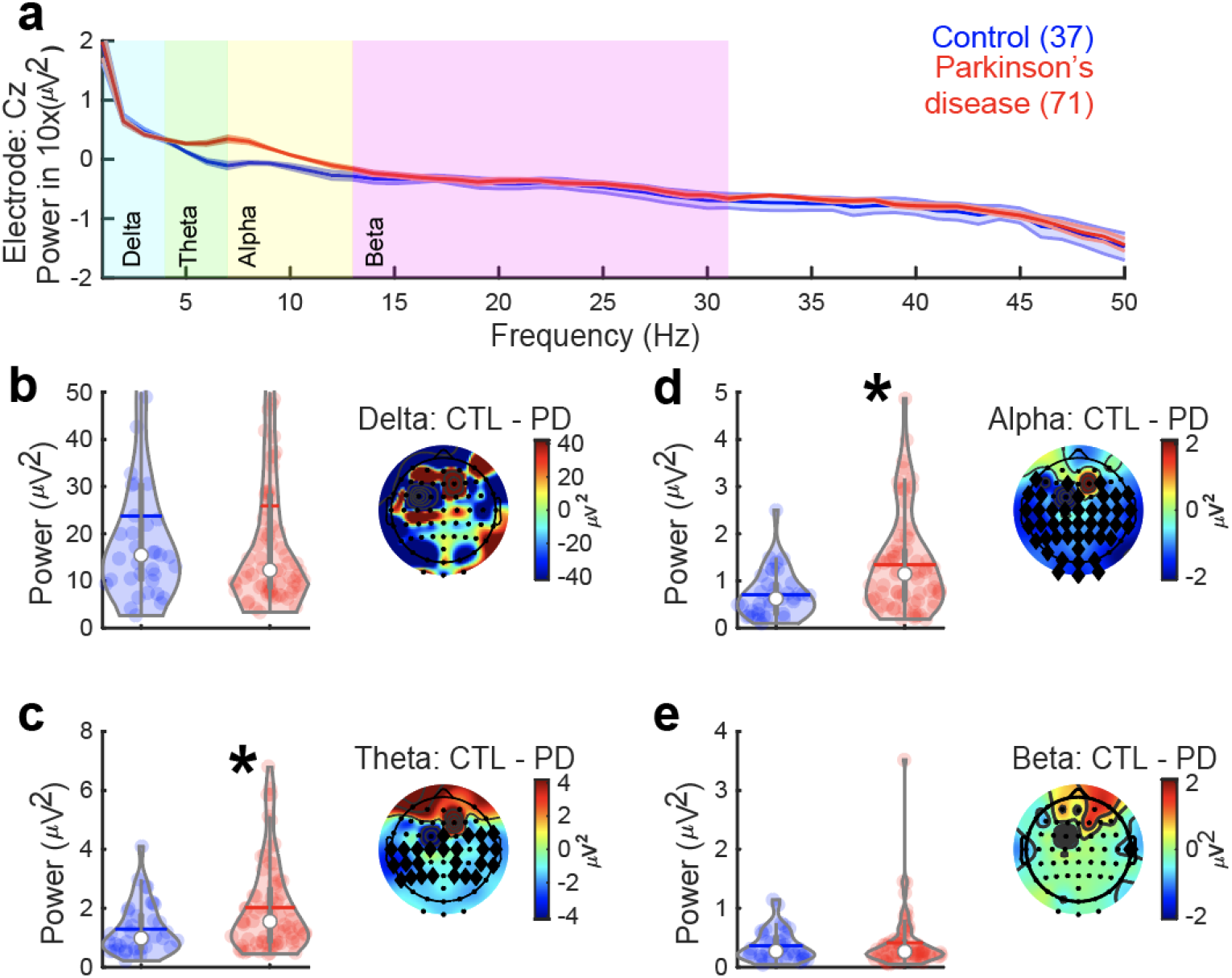
Baseline midfrontal theta and alpha power is increased in PD. We also performed conventional spectral analysis on pre-stimulation data following Welch’s power spectral density estimation method, using the pwelch function in MATLAB. We selected a 1-second time window prior to the instructional cue. The number of overlapped samples was set to 50% of the window length and the non-equispaced fast Fourier transform (NFFT) was assigned as the length of the segments. a) Resting-state power spectral density of EEG power at midfrontal electrode Cz revealed marked increases in theta (4-7 Hz) and alpha (7-13 Hz) power for control (blue) and PD patients (red). Resting-state epoch was 1 second prior to the instructional text from 7-second interval trials. Power for control and PD patients for b) delta bands (1-4 Hz); c) theta bands (4-7 Hz; * = t_(106)_=-2.70, p=0.008); d) alpha bands (7-13 Hz; * = alpha: t_(106)_=-3.56, p=0.001); and e) beta bands (13-30 Hz).

**Fig. S4.**
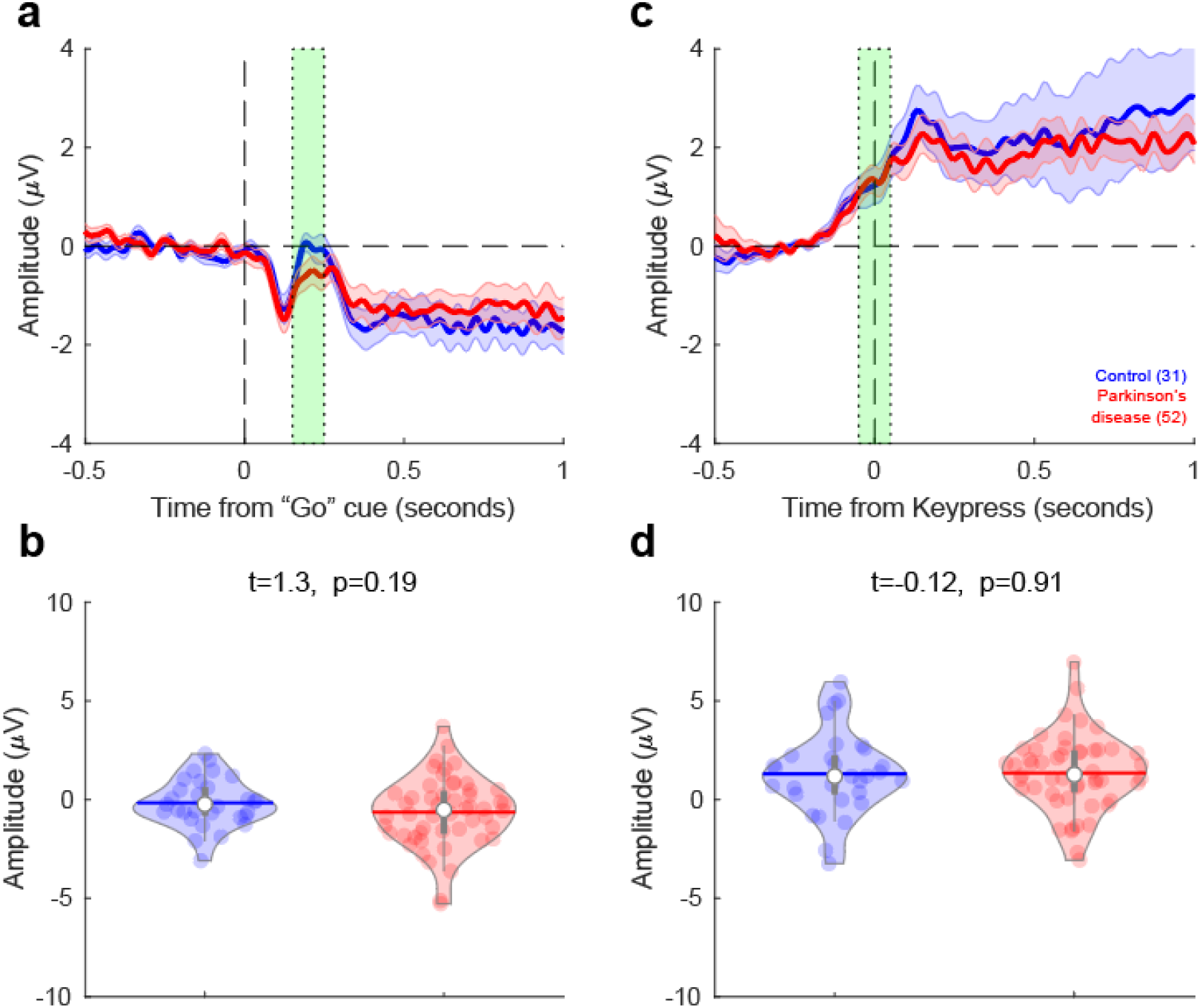
Event-related potentials (ERP) from electrode Cz for 3-second trials. No difference in midfrontal cue-triggered (a and b) and response-triggered (c and d) ERPs during 3-second interval timing task was observed between PD patients and control subjects (ROIs: green box).

**Fig. S5.**
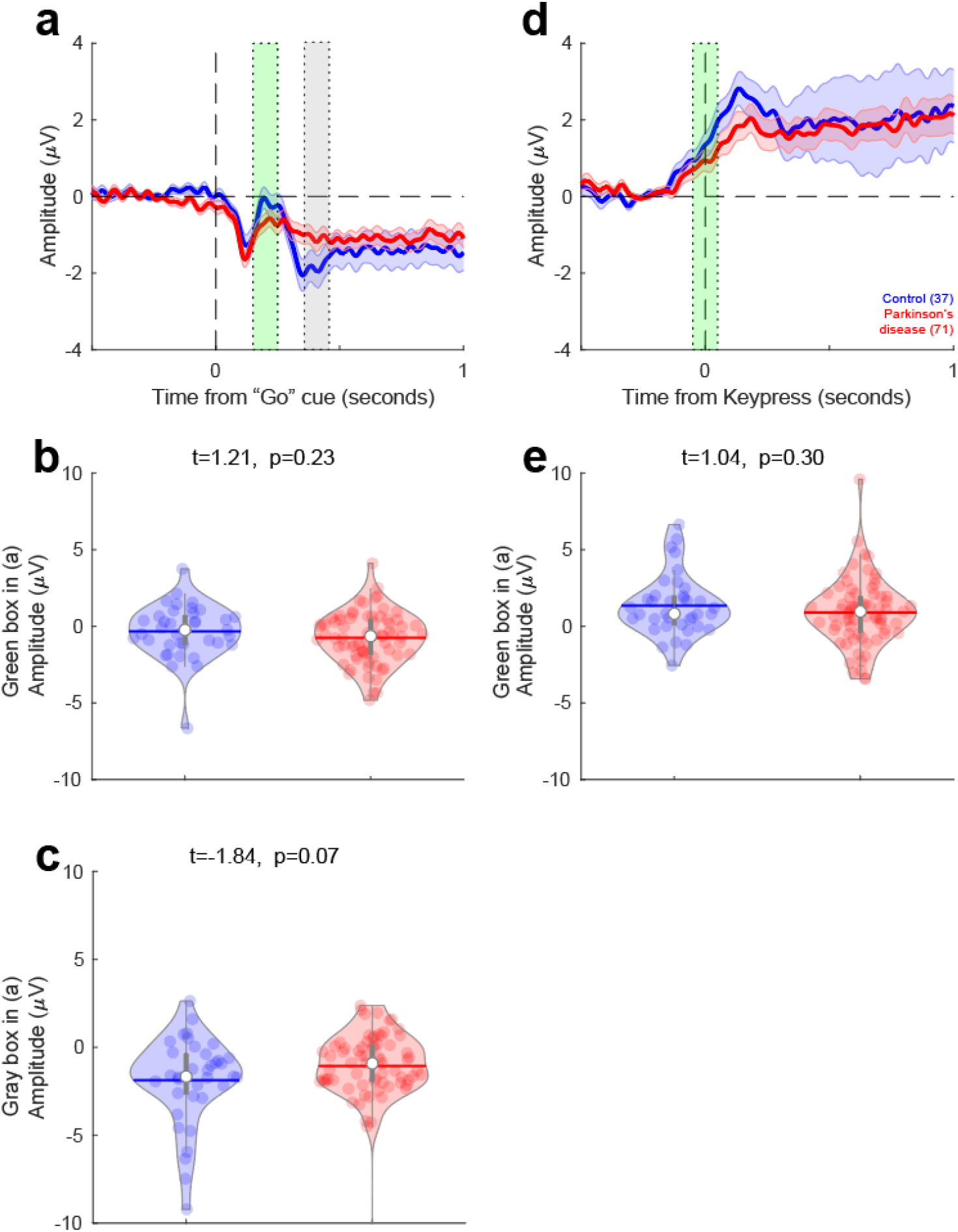
Event-related potentials (ERP) from electrode Cz for 7-second trials. No reliable differences in midfrontal cue-triggered (a, b, c, and d-e) and keypress-triggered ERPs during 7-second interval timing task was observed between PD patients and control subjects (ROIs: green/gray box for cue-triggered analysis, green-box for keypress-triggered analysis).

**Fig. S6.**
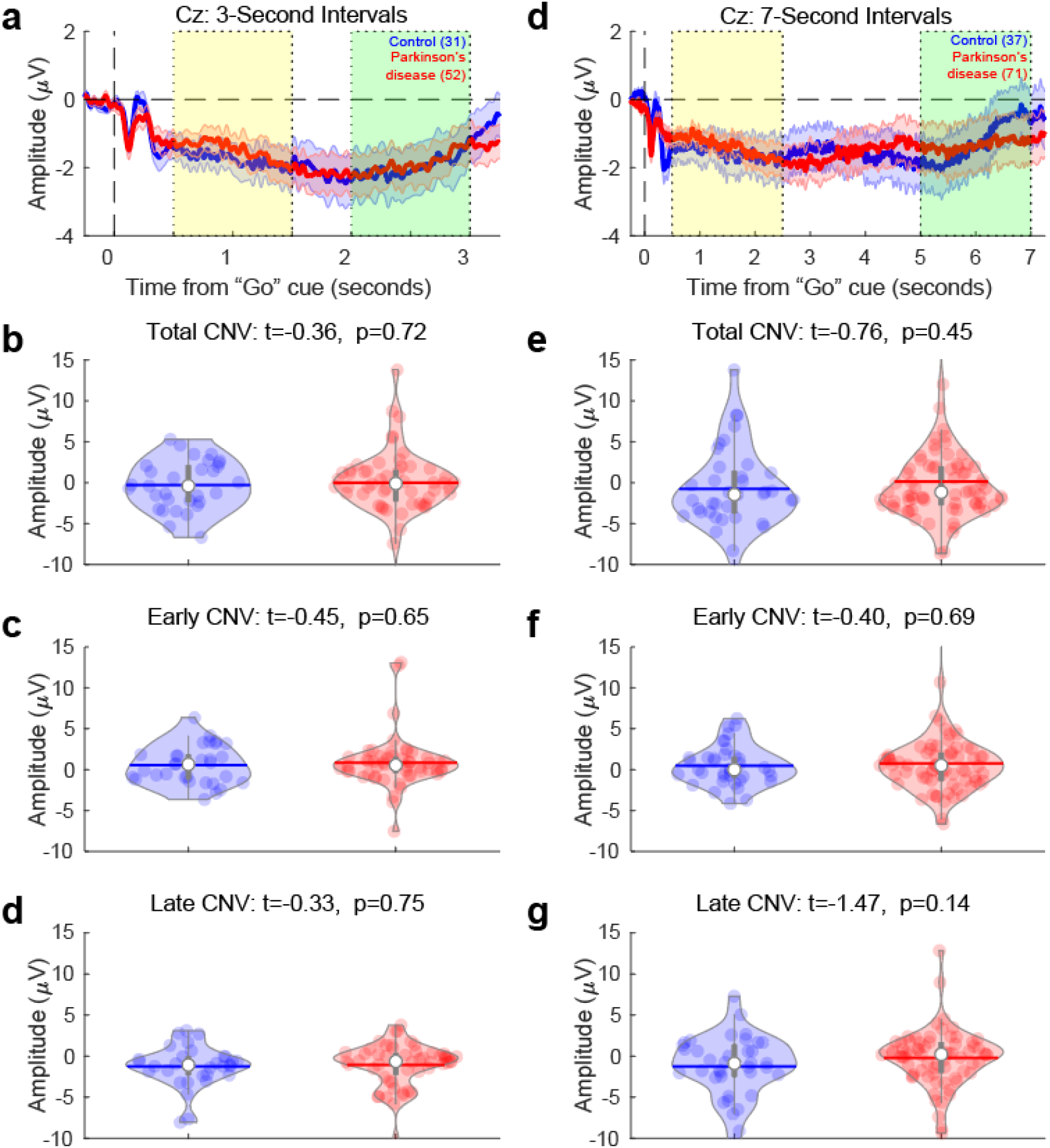
Contingent-negative variation (CNV) analyses for early and late epochs. For 3-second intervals, a) we found no reliable differences in midfrontal CNV activity between control and PD participants for the mean amplitude of b) total CNV (0.5-3 seconds), c) early CNV (0.5 – 1.5 seconds), or d) late CNVs (2-3 seconds). We also observed no reliable differences for e) 7-second intervals for f) total CNV (0.5-7 seconds), g) early CNV (0.5-2.5 seconds), or h) late CNVs (5-7 seconds).

**Fig. S7.**
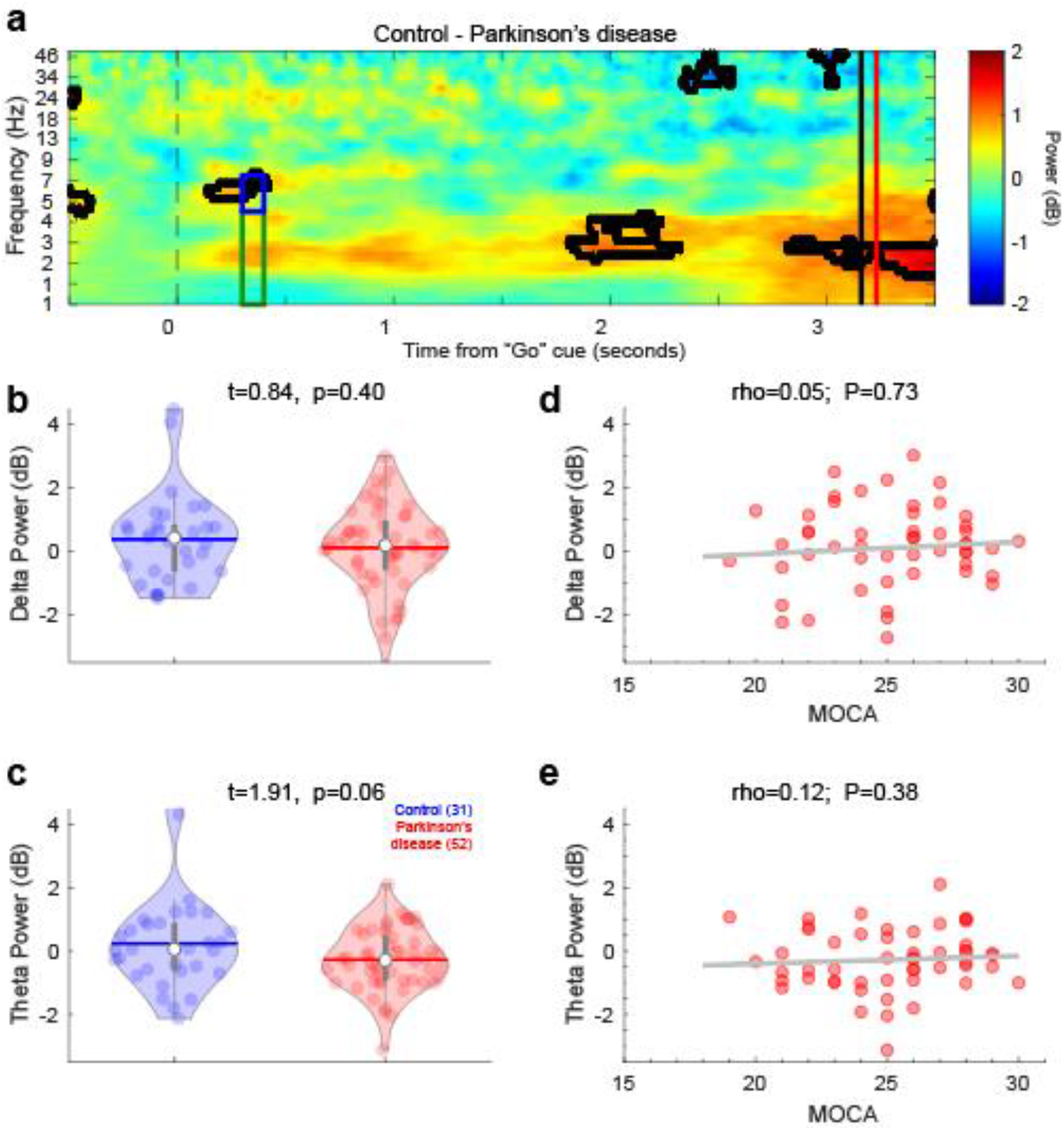
Midfrontal delta/theta activity for 3-second intervals. a) We compared time-frequency power of midfrontal activity after the imperative “Go” cue for 3-second trials. Solid black lines indicate p<0.05 via a t-test of activity in control compared to PD patients. b) cue-triggered midfrontal delta power (1-4 Hz, tf-ROI: green box in a) and c) cue-triggered midfrontal theta power (4-7 Hz, tf-ROI: blue box in a) in PD patients. d) delta and e) theta power vs. cognitive dysfunction as measured by MOCA in PD patients. Data from PD (n=52) and control (n=31) patients on 3-second trials.

**Fig. S8.**
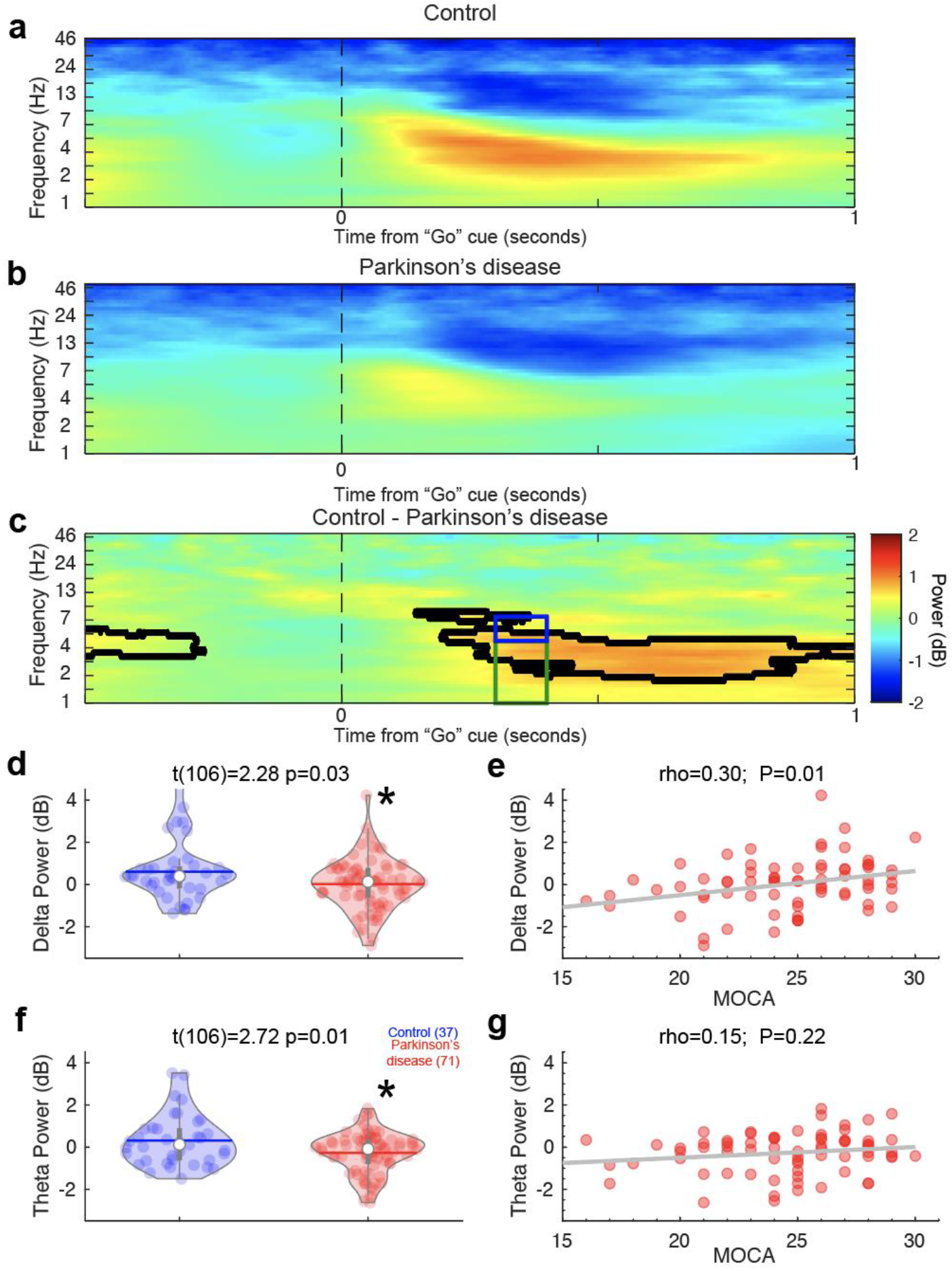
Cue-triggered delta power from a cluster of midfrontal electrodes (FCz, Cz, and CPz) predicts cognitive dysfunction in PD. Whereas data in Figure 2 included electrode Cz according to our *a priori* hypothesis, data in this figure includes an expanded midfrontal coverage including FCz, Cz, CPz. Frequency power of midfrontal activity over time from the imperative “Go” cue from a) control and b) PD participants on 7-second trials. c) Comparison of control and PD patients. Areas outlined by solid black lines indicate p<0.05 via a t-test of activity in control compared to PD participants. There was significantly less d) cue-triggered midfrontal delta power (1-4 Hz, time-frequency-Region-of-interest (tf-ROI): green box) and f) cue-triggered midfrontal theta power (4-7 Hz, tf-ROI: blue box) in controls vs. PD patients. e) Delta power predicted cognitive dysfunction as measured by MOCA in PD patients, but g) theta power did not.*=p<0.05. Data from control (n=37) and PD (n=71) patients.

**Fig. S9.**
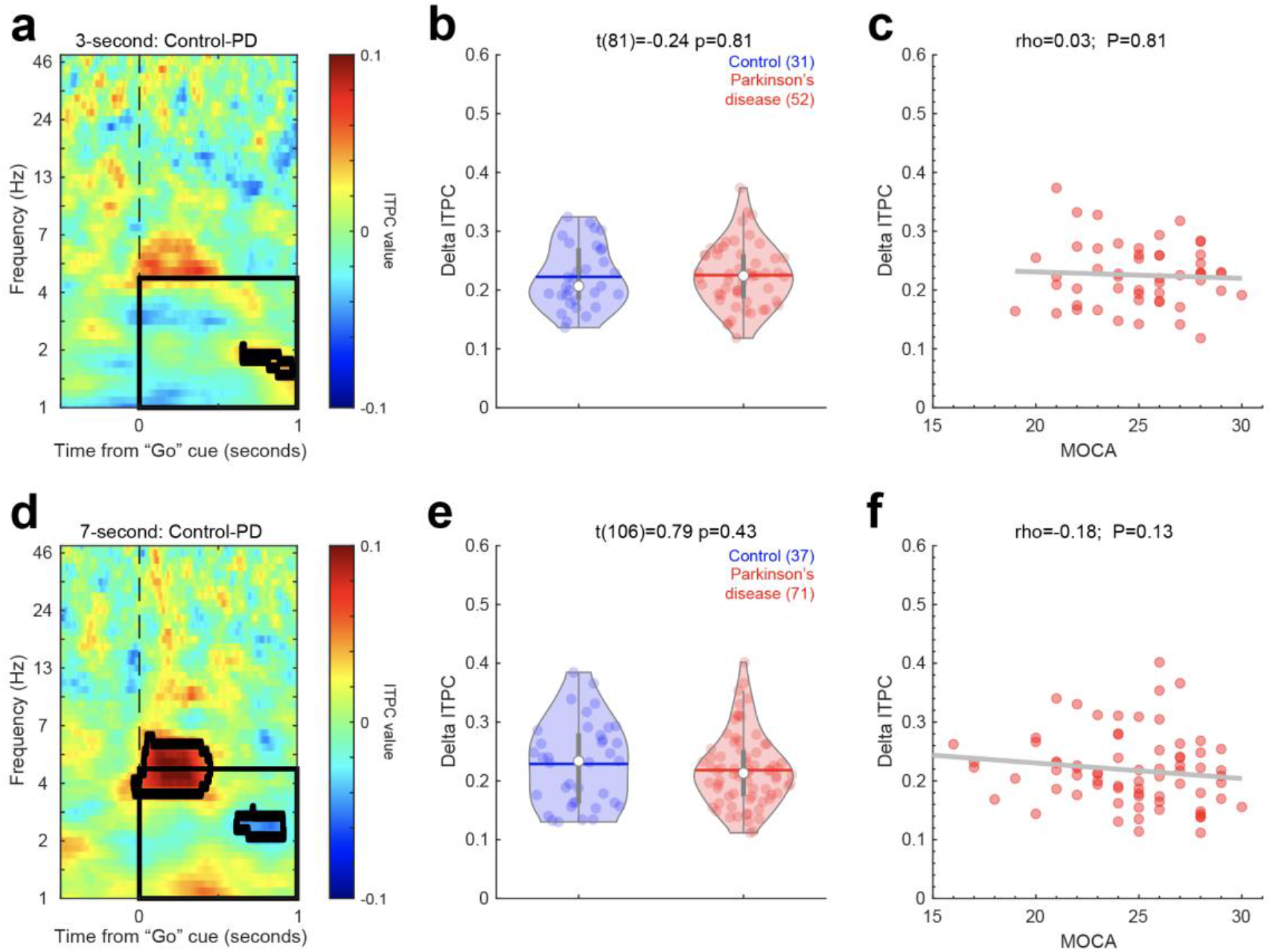
Intertrial phase coherence (ITPC). a) For 3-second trials, there were, b) few overall consistent differences between Control and PD patients for delta bands 0-1 seconds after the cue, and c) no relationship with MOCA. Data from 24 ± 1.4 (mean ± SEM) trials. d) For 7-second trials, there were marked differences in ~4 Hz ITPC around the go-cue, but e) no overall differences in delta bands 0-1 seconds after the cue, and f) no relationship with MOCA. Data from 33 ± 0.8 trials.

**Fig. S10.**
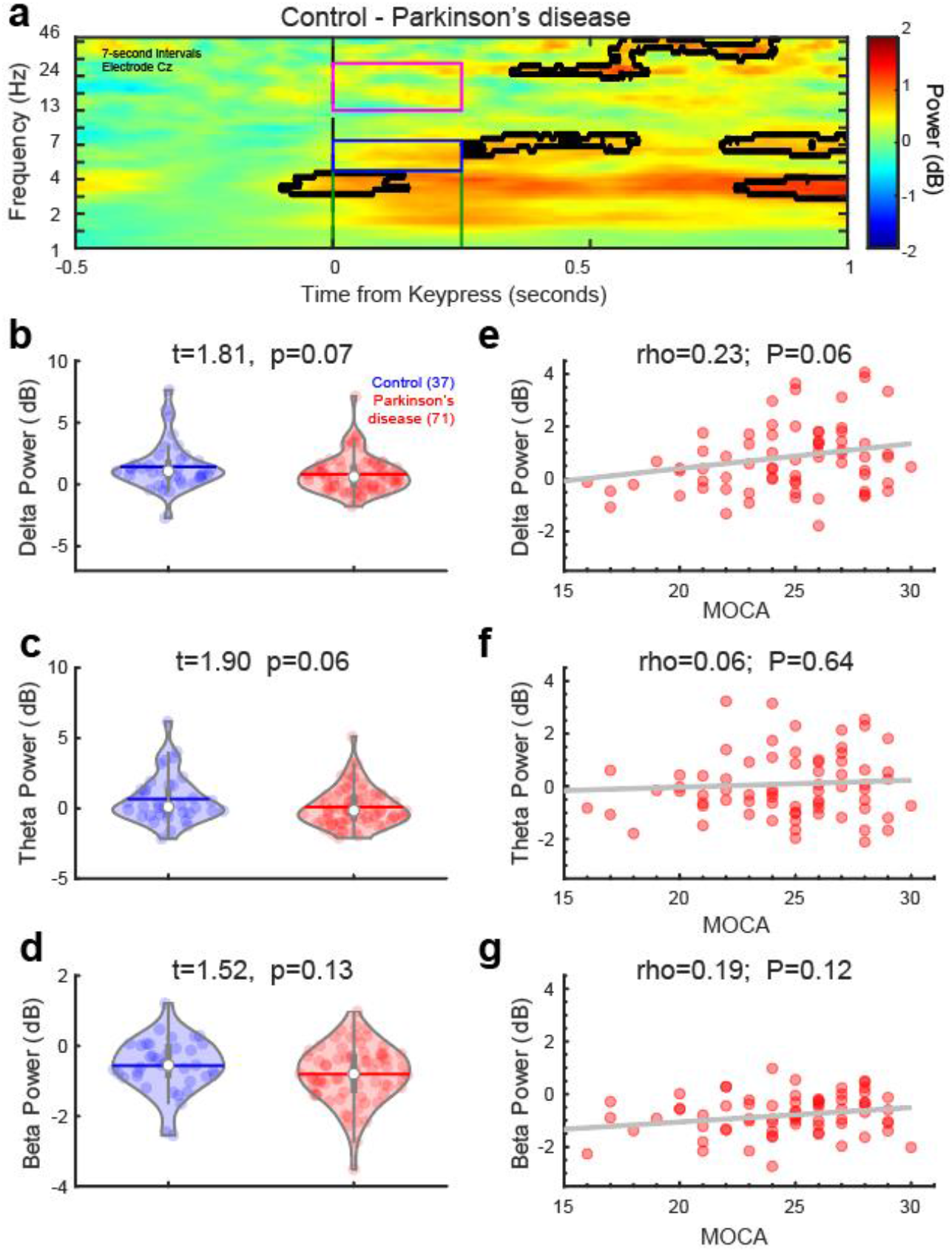
Midfrontal activity around keypress. a) We compared time-frequency power of midfrontal activity around response (“keypress”; time=0). Areas outlined by solid black lines indicate p<0.05 via a t-test of activity in control compared to PD participants. b) delta power (1-4 Hz, tf-ROI: green box in a), c) theta power (4-7 Hz, tf-ROI: blue box in a) in PD patients, and d) beta power (13-30 Hz, tf-ROI: magenta box in a) in control vs. PD patients. e) delta, f) theta, and g) beta power do not significantly correlate with cognitive dysfunction as measured by MOCA in PD patients. Data from 71 PD and 37 control patients on 7-second trials.

**Fig. S11.**
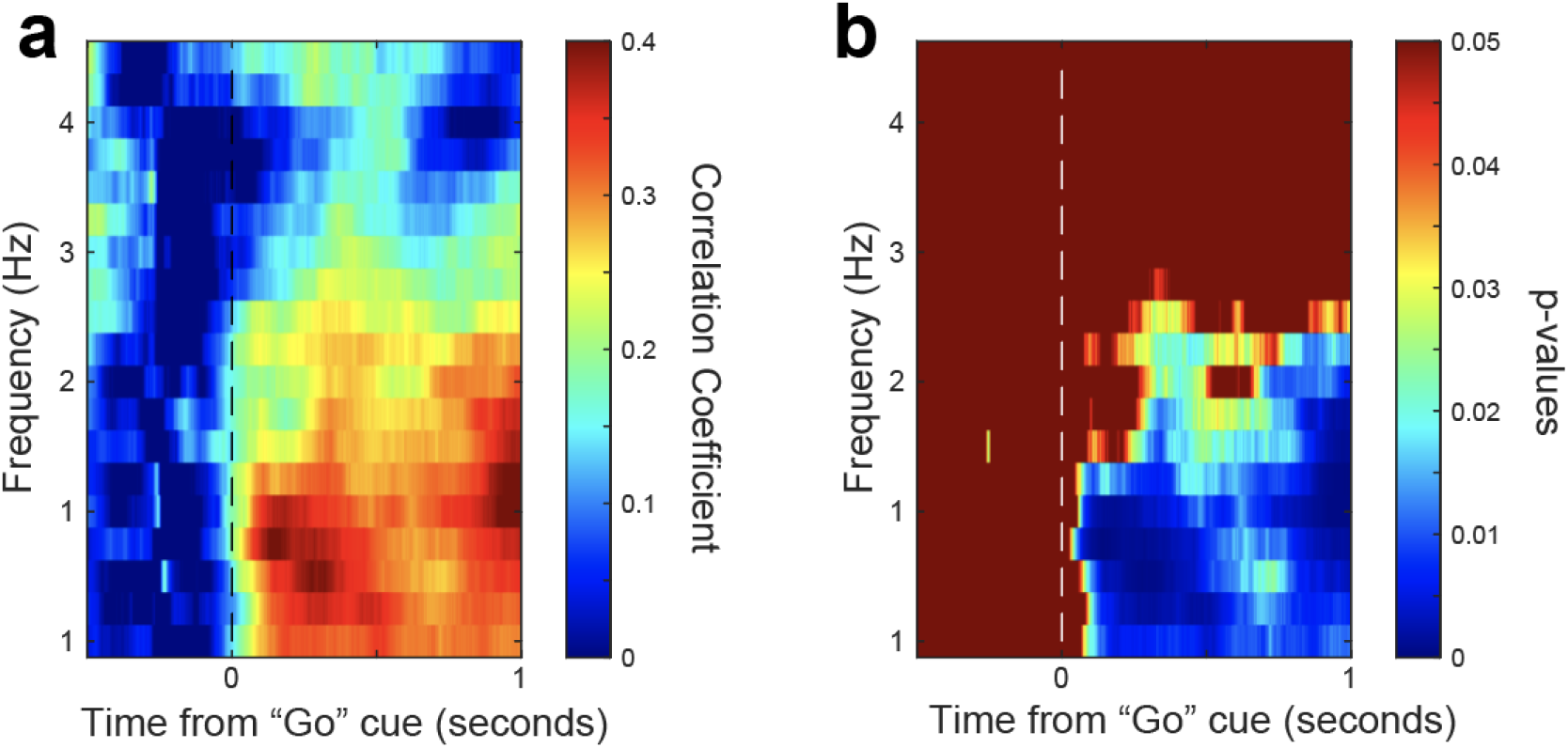
Correlations of cue-triggered delta power with MOCA. a). Correlation coefficient, and b) p-values for each point on time-frequency spectrogram from EEG electrode Cz. Data from 71 PD patients in Figure 2.

**Fig. S12.**
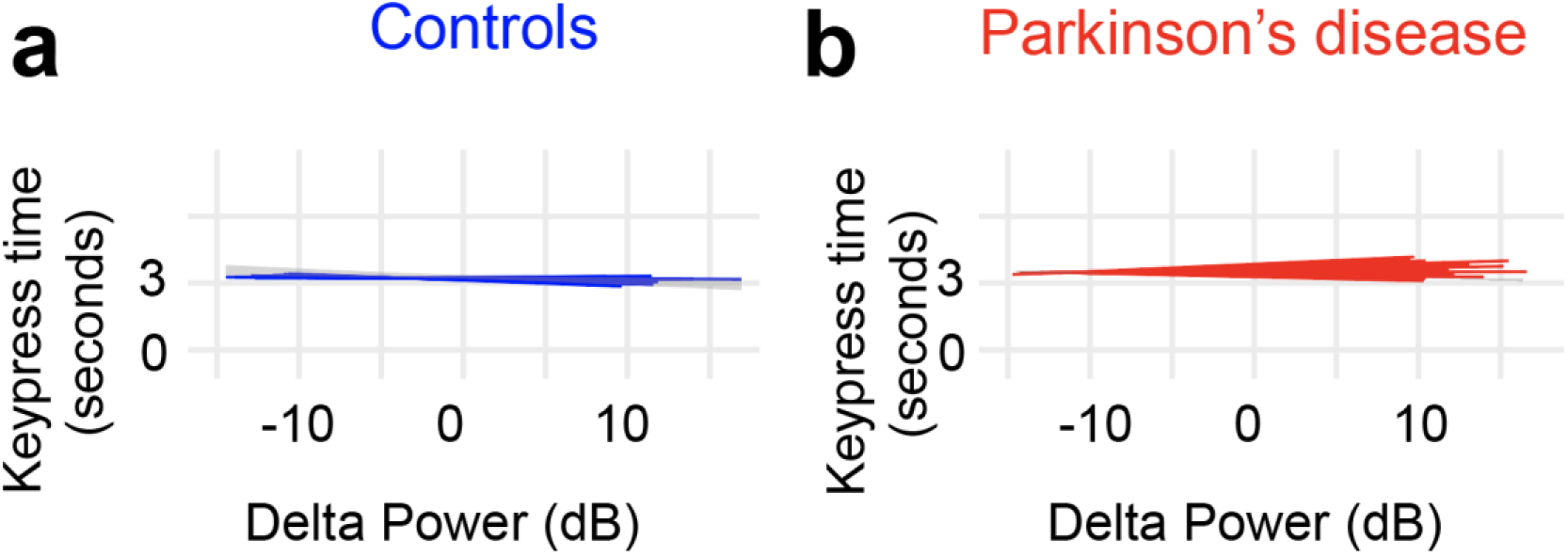
Delta activity does not predict temporal variability for 3-second intervals. a-b) No clear relationship with delta power and time estimates was observed in controls or PD patients during 3-second interval timing task.

**Fig. S13.**
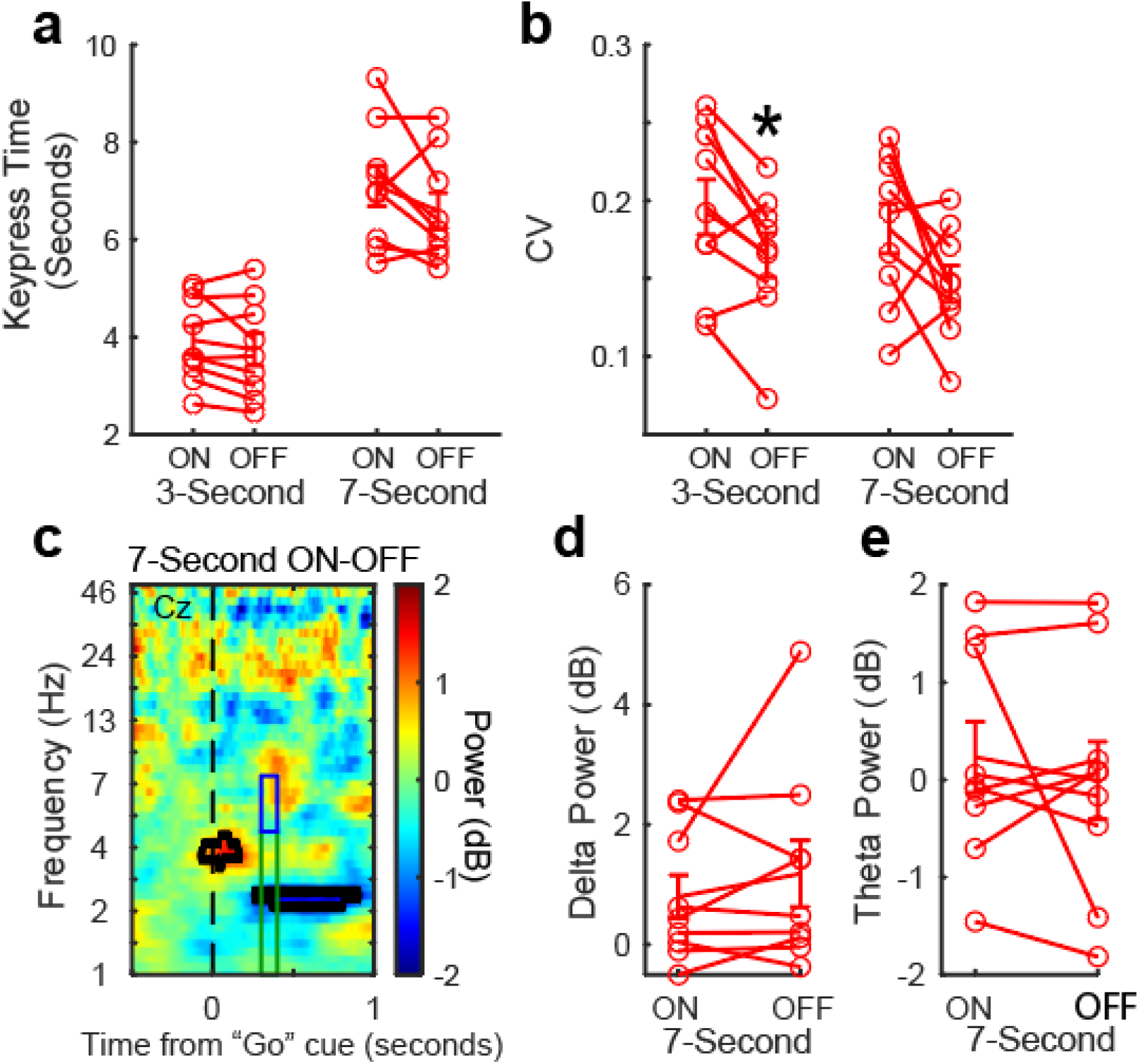
Levodopa does not reliably affect midfrontal delta-theta activity. In PD patients, we examined a) mean response time, b) keypress CV, c) midfrontal cue-triggered activity from PD patients ON versus OFF levodopa. Areas outlined by solid black lines indicate p<0.05 via a t-test of activity in control compared to PD participants. d) Midfrontal cue-triggered delta activity (tfROI: green box) in c) and e) midfrontal cue-triggered theta-activity (tfROI: blue box in c) from PD patients ON and OFF levodopa. Data from 9 patients; *=p<0.05 via paired t-test.

## Notes

### Competing Interest Statement

The authors have declared no competing interest.

## REFERENCES

1. Aarsland, D., Brønnick, K., Larsen, J.P., Tysnes, O.B. & Alves, G. Cognitive impairment in incident, untreated Parkinson disease: the Norwegian ParkWest study. Neurology 72, 1121–1126 (2009).

2. Alberico, S.L., Cassell, M.D. & Narayanan, N.S. The Vulnerable Ventral Tegmental Area in Parkinson’s Disease. Basal Ganglia 5, 51–55 (2015).

3. Hely, M.A., Reid, W.G.J., Adena, M.A., Halliday, G.M. & Morris, J.G.L. The Sydney multicenter study of Parkinson’s disease: the inevitability of dementia at 20 years. Mov. Disord. 23, 837–844 (2008).

4. Williams-Gray, C.H., Foltynie, T., Brayne, C.E.G., Robbins, T.W. & Barker, R.A. Evolution of cognitive dysfunction in an incident Parkinson’s disease cohort. Brain 130, 1787–1798 (2007).

5. Deuschl, G. et al. A randomized trial of deep-brain stimulation for Parkinson’s disease. N. Engl. J. Med. 355, 896–908 (2006).

6. Follett, K.A. Comparison of pallidal and subthalamic deep brain stimulation for the treatment of levodopa-induced dyskinesias. Neurosurg. Focus 17(2004).

7. Limousin, P. et al. Electrical stimulation of the subthalamic nucleus in advanced Parkinson’s disease. N. Engl. J. Med. 339, 1105–1111 (1998).

8. Little, S. et al. Adaptive deep brain stimulation in advanced Parkinson disease. Ann. Neurol. 74, 449–457 (2013).

9. Chaudhuri, K.R. & Odin, P. The challenge of non-motor symptoms in Parkinson’s disease. Prog. Brain Res. 184, 325–341 (2010).

10. Forsaa, E.B., Larsen, J.P., Wentzel-Larsen, T. & Alves, G. What predicts mortality in Parkinson disease?: a prospective population-based long-term study. Neurology 75, 1270–1276 (2010).

11. Aldridge, G.M., Birnschein, A., Denburg, N.L. & Narayanan, N.S. Parkinson’s Disease Dementia and Dementia with Lewy Bodies Have Similar Neuropsychological Profiles. Front. Neurol. 9, 123 (2018).

12. Owen, A.M. Cognitive dysfunction in Parkinson’s disease: the role of frontostriatal circuitry. Neuroscientist 10, 525–537 (2004).

13. Buhusi, C.V. & Meck, W.H. What makes us tick? Functional and neural mechanisms of interval timing. Nat. Rev. Neurosci. 6, 755–765 (2005).

14. Brück, A. et al. Positron emission tomography shows that impaired frontal lobe functioning in Parkinson’s disease is related to dopaminergic hypofunction in the caudate nucleus. Neurosci. Lett. 311, 81–84 (2001).

15. Coull, J.T., Cheng, R.-K. & Meck, W.H. Neuroanatomical and neurochemical substrates of timing. Neuropsychopharmacology 36, 3–25 (2011).

16. Parker, K.L., Lamichhane, D., Caetano, M.S. & Narayanan, N.S. Executive dysfunction in Parkinson’s disease and timing deficits. Front. Integr. Neurosci. 7, 75 (2013).

17. Malapani, C., Deweer, B. & Gibbon, J. Separating storage from retrieval dysfunction of temporal memory in Parkinson’s disease. J. Cogn. Neurosci. 14, 311–322 (2002).

18. Malapani, C. et al. Coupled temporal memories in Parkinson’s disease: a dopamine-related dysfunction. J. Cogn. Neurosci. 10, 316–331 (1998).

19. Jones, C.R. & Jahanshahi, M. Contributions of the basal ganglia to temporal processing: Evidence from Parkinson’s disease. Timing and Time Perception 1, 1–41 (2013).

20. Merchant, H., Luciana, M., Hooper, C., Majestic, S. & Tuite, P. Interval timing and Parkinson’s disease: heterogeneity in temporal performance. Exp. Brain Res. 184, 233–248 (2008).

21. Kim, Y.-C. et al. Optogenetic Stimulation of Frontal D1 Neurons Compensates for Impaired Temporal Control of Action in Dopamine-Depleted Mice. Curr. Biol. 27, 39–47 (2017).

22. Parker, K.L., Chen, K.-H., Kingyon, J.R., Cavanagh, J.F. & Narayanan, N.S. Medial frontal ∼4-Hz activity in humans and rodents is attenuated in PD patients and in rodents with cortical dopamine depletion. J. Neurophysiol. 114, 1310–1320 (2015).

23. Cavanagh, J.F. & Frank, M.J. Frontal theta as a mechanism for cognitive control. Trends Cogn. Sci. 18, 414–421 (2014).

24. Cavanagh, J.F., Zambrano-Vazquez, L. & Allen, J.J.B. Theta lingua franca: A common mid-frontal substrate for action monitoring processes. Psychophysiology 49, 220–238 (2012).

25. Narayanan, N.S., Cavanagh, J.F., Frank, M.J. & Laubach, M. Common medial frontal mechanisms of adaptive control in humans and rodents. Nat. Neurosci. 16, 1888–1897 (2013).

26. Cosman, J.D., Lowe, K.A., Zinke, W., Woodman, G.F. & Schall, J.D. Prefrontal Control of Visual Distraction. Curr. Biol. 28, 414–420.e413 (2018).

27. Chen, K.-H. et al. Startle Habituation and Midfrontal Theta Activity in Parkinson’s Disease. J. Cogn. Neurosci., 1–11 (2016).

28. Singh, A., Richardson, S.P., Narayanan, N. & Cavanagh, J.F. Mid-frontal theta activity is diminished during cognitive control in Parkinson’s disease. Neuropsychologia 117, 113–122 (2018).

29. Kelley, R. et al. A human prefrontal-subthalamic circuit for cognitive control. Brain 141, 205–216 (2018).

30. Wojtecki, L. et al. Modulation of Human Time Processing by Subthalamic Deep Brain Stimulation. PLoS One 6(2011).

31. Church, R.M. Properties of the internal clock. Ann. N. Y. Acad. Sci. 423, 566–582 (1984).

32. Gibbon, J., Church, R.M. & Meck, W.H. Scalar timing in memory. Ann. N. Y. Acad. Sci. 423, 52–77 (1984).

33. Singh, A. et al. Frontal theta and beta oscillations during lower-limb movement in Parkinson’s disease. Clin. Neurophysiol. 131, 694–702 (2020).

34. Martinu, K., Degroot, C., Madjar, C., Strafella, A.P. & Monchi, O. Levodopa influences striatal activity but does not affect cortical hyper-activity in Parkinson’s disease. Eur. J. Neurosci. 35, 572–583 (2012).

35. Narayanan, N.S., Rodnitzky, R.L. & Uc, E.Y. Prefrontal dopamine signaling and cognitive symptoms of Parkinson’s disease. Rev. Neurosci. 24, 267–278 (2013).

36. Wingeier, B. et al. Intra-operative STN DBS attenuates the prominent beta rhythm in the STN in Parkinson’s disease. Exp. Neurol. 197, 244–251 (2006).

37. Okun, M.S. Deep-brain stimulation for Parkinson’s disease. N. Engl. J. Med. 367, 1529–1538 (2012).

38. Singh, A. Oscillatory activity in the cortico-basal ganglia-thalamic neural circuits in Parkinson’s disease. Eur. J. Neurosci. 48, 2869–2878 (2018).

39. Wearden, J.H. et al. Stimulus timing by people with Parkinson’s disease. Brain Cogn. 67, 264–279 (2008).

40. Jensen, O. & Tesche, C.D. Frontal theta activity in humans increases with memory load in a working memory task. Eur. J. Neurosci. 15, 1395–1399 (2002).

41. Kim, Y.-C. & Narayanan, N.S. Prefrontal D1 Dopamine-Receptor Neurons and Delta Resonance in Interval Timing. Cereb. Cortex (2018).

42. Parker, K.L., Chen, K.-H., Kingyon, J.R., Cavanagh, J.F. & Narayanan, N.S. D1-Dependent 4 Hz Oscillations and Ramping Activity in Rodent Medial Frontal Cortex during Interval Timing. J. Neurosci. 34, 16774–16783 (2014).

43. Herz, D.M., Zavala, B.A., Bogacz, R. & Brown, P. Neural Correlates of Decision Thresholds in the Human Subthalamic Nucleus. Curr. Biol. 26, 916–920 (2016).

44. Fujisawa, S. & Buzsáki, G. A 4 Hz oscillation adaptively synchronizes prefrontal, VTA, and hippocampal activities. Neuron 72, 153–165 (2011).

45. Emmons, E.B. et al. Rodent Medial Frontal Control of Temporal Processing in the Dorsomedial Striatum. J. Neurosci. 37, 8718–8733 (2017).

46. Wang, J., Narain, D., Hosseini, E.A. & Jazayeri, M. Flexible timing by temporal scaling of cortical responses. Nat. Neurosci. 21, 102 (2018).

47. Paton, J.J. & Buonomano, D.V. The Neural Basis of Timing: Distributed Mechanisms for Diverse Functions. Neuron 98, 687–705 (2018).

48. Scangos, K.W., Carter, C.S., Gurkoff, G., Zhang, L. & Shahlaie, K. A pilot study of subthalamic theta frequency deep brain stimulation for cognitive dysfunction in Parkinson’s disease. Brain Stimul. 11, 456–458 (2018).

49. Parker, K.L. et al. Delta-frequency stimulation of cerebellar projections can compensate for schizophrenia-related medial frontal dysfunction. Mol. Psychiatry 22, 647–655 (2017).

50. Aarsland, D. & Kurz, M.W. The Epidemiology of Dementia Associated with Parkinson’s Disease. Brain Pathol. 20, 633–639 (2010).

51. Morrison, C.E., Borod, J.C., Brin, M.F., Hälbig, T.D. & Olanow, C.W. Effects of levodopa on cognitive functioning in moderate-to-severe Parkinson’s disease (MSPD). J. Neural Transm. 111, 1333–1341 (2004).

52. Cools, R. Dopaminergic modulation of cognitive function-implications for L-DOPA treatment in Parkinson’s disease. Neurosci. Biobehav. Rev. 30, 1–23 (2006).

53. Cools, R. & D’Esposito, M. Inverted-U-shaped dopamine actions on human working memory and cognitive control. Biol. Psychiatry 69, e113–125 (2011).

54. Cavanagh, J.F., Kumar, P., Mueller, A.A., Richardson, S.P. & Mueen, A. Diminished EEG habituation to novel events effectively classifies Parkinson’s patients. Clin. Neurophysiol. 129, 409–418 (2018).

55. Brown, D.R., Richardson, S.P. & Cavanagh, J.F. An EEG marker of reward processing is diminished in Parkinson’s disease. Brain Res. 1727, 146541 (2020).

56. Rakitin, B.C., Scarmeas, N., Li, T., Malapani, C. & Stern, Y. Single-dose levodopa administration and aging independently disrupt time production. J. Cogn. Neurosci. 18, 376–387 (2006).

57. Rakitin, B.C. et al. Scalar expectancy theory and peak-interval timing in humans. J. Exp. Psychol. 24, 15–33 (1998).

58. Delorme, A. & Makeig, S. EEGLAB: an open source toolbox for analysis of single-trial EEG dynamics including independent component analysis. J. Neurosci. Methods 134, 9–21 (2004).

59. Seer, C., Lange, F., Georgiev, D., Jahanshahi, M. & Kopp, B. Event-related potentials and cognition in Parkinson’s disease: An integrative review. Neurosci. Biobehav. Rev. 71, 691–714 (2016).

60. Linssen, A.M. et al. Contingent negative variation as a dopaminergic biomarker: evidence from dose-related effects of methylphenidate. Psychopharmacology (Berl.) 218, 533–542 (2011).

61. Cohen, M.X. Analyzing Neural Time Series Data: Theory and Practice (Issues in Clinical and Cognitive Neuropsychology). (2014).

62. Biundo, R. et al. MMSE and MoCA in Parkinson’s disease and dementia with Lewy bodies: a multicenter 1-year follow-up study. J. Neural Transm. 123, 431–438 (2016).

63. Dalrymple-Alford, J.C. et al. The MoCA: well-suited screen for cognitive impairment in Parkinson disease. Neurology 75, 1717–1725 (2010).

64. Movement Disorder Society Task Force on Rating Scales for Parkinson’s, D. The Unified Parkinson’s Disease Rating Scale (UPDRS): status and recommendations. Mov. Disord. 18, 738–750 (2003).

65. Cavanagh, J.F., Napolitano, A., Wu, C. & Mueen, A. The Patient Repository for EEG Data + Computational Tools (PRED+CT). Front. Neuroinform. 11, 67 (2017).

